# A new single-cell level R-index for EGFR-TKI resistance and survival prediction in LUAD

**DOI:** 10.1101/2021.07.30.454426

**Authors:** Xiaohong Xie, Lifeng Li, Liang Xie, Zhentian Liu, Xuan Gao, Xuefeng Xia, Haiyi Deng, Yilin Yang, MeiLing Yang, Lianpeng Chang, Xin Yi, Zhiyi He, Chengzhi Zhou

**Affiliations:** State Key Laboratory of Respiratory Disease, National Clinical Research Centre for Respiratory Disease, Guangzhou Institute of Respiratory Health, The First Affiliated Hospital of Guangzhou Medical University, Guangzhou Medical University, Guangzhou, Guangdong, 510120, China; Department of Pulmonary and Critical Care Medicine, The First Affiliated Hospital of Guangxi Medical University, Nanning, Guangxi, 530021, China; Department of Thoracic Surgery, Guangdong Provincial People’s Hospital / Guangdong Academy of Medical Sciences, Guangzhou, Guangdong, 510080, China; Geneplus-Beijing, Beijing, 102206, China; State Key Laboratory of Microbial Resources, Institute of Microbiology, Chinese Academy of Sciences, Beijing, 100101, China; Geneplus- Shenzhen Clinical Laboratory, Shenzhen, Guangdong, 518122, China

**Author notes:** Corresponding authors: Prof. Chengzhi Zhou, State Key Laboratory of Respiratory Disease, National Clinical Research Centre for Respiratory Disease, Guangzhou Institute of Respiratory Health, The First Affiliated Hospital of Guangzhou Medical University, Guangzhou Medical University, Guangzhou, Guangdong, 510120, China, Prof. Zhiyi He, Department of Pulmonary and Critical Care Medicine, The First Affiliated Hospital of Guangxi Medical University, Nanning, Guangxi 530021, China. Joint Authors contributed equally (Xiaohong Xie, Lifeng Li, Liang Xie).

**Keywords:** EGFR-TKI resistance, lung adenocarcinoma, R-index/scRNA-seq

## Abstract

EGFR-TKIs achieved excellent efficacy in EGFR-mutated patients. Unfortunately, most patients would inevitably develop progressive disease within a median of 10 to 14 months. Predicting the resistance probability remains a challenge. Therefore, we created an R-index model trained by single-cell RNA data with the OCLR algorithm. This model can be applied to estimate the level of EGFR-TKI resistance in cell line and xenograft mice models and predict prognosis in multiple cohorts. Comparing the high and the low R-index group, we found that the glycolysis pathway and KRAS up-regulation pathway were related to resistance, and MDSC was the leading cause of immunosuppression in the tumor microenvironment. These results are consistent with previous studies indicating that the R-index provides an insight into resistance status and a new way to explore resistance mechanisms and clinical treatment by the combination of Glucose metabolism-targeted or MDSC-targeted therapies. This is the first quantification method of EGFR-TKI resistance based on single-cell sequencing data solving the problem of the mixed resistance state of tumor cells and helping explore transcriptome characteristics of drug-resistant cell populations.

## INTRODUCTION

Lung cancer is the leading cause of cancer death worldwide(Siegel, Miller et al., 2021). According to previous reports, approximately 85% of patients are diagnosed with non-small cell lung cancer (NSCLC), of which lung adenocarcinoma (LUAD) was the most common histological subtype (Molina, Yang et al., 2008). With the development of NGS technology, the treatment of LUAD has developed from the empirical use of radiotherapy and chemotherapy to various targeted therapies and immunotherapy(2014). And these increasingly comprehensive therapies are still needed to improve patient’s clinical outcomes and quality of life. There are two enlightening treatment concepts for patients with advanced lung adenocarcinoma: extinction therapy and adaptive therapy(Gatenby & Brown, 2020). This extinction therapy scheme is widely used in clinical practice, combined with imaging information and patient status to formulate a multi-line treatment plan. When therapeutic implications were found in tumor tissue, initial treatment (Single drug or multi-drug combination, first strike) is used to create an extinction threshold limiting and destroying the tumor cells, and regrow resistance is subsequently treated with new drugs (second strike) (Johnson, Howard et al., 2019, Walther, Hiley et al., 2015, Wu & Shih, 2018). The strategy of adaptive therapy is to utilize intratumoral evolutionary dynamics to suppress the proliferation of resistant tumor cells and prolong the response to treatment. Drug-sensitive tumor cells could compete with the drug-resistant tumor cells without drug interference, and give the time window allowing for the competition would restrain the proliferation of tolerant cells. Although evidence in lung cancer showed that after a drug holiday, EGFR-TKI resistance patients regain the sensitivity have proven it to be a feasible method, the design of adaptive therapy is still challenging(Ohashi, Maruvka et al., 2013, Yamaguchi, Kaira et al., 2019).

Epidermal growth factor receptor tyrosine kinase inhibitors (EGFR-TKIs) are used in the targeted therapy for LUAD patients with oncogenic drivers of EGFR mutations since 2004 and have reported a high response rate of 80% (Lynch, Bell et al., 2004, Paez, Jänne et al., 2004). The most common type of acquired resistance to the first and second generation of EGFR-TKIs is caused by the secondary mutation EGFR T790M (Arcila, Oxnard et al., 2011). Although the third-generation EGFR-TKIs have been clinically available targeting EGFR T790M, most patients progress within a median of 10 to 14 months of treatment (Rosell, Moran et al., 2009). The resistance mechanisms include PIK3CA mutations, BRAF mutations, c-MET amplification, AXL overexpression, small-cell lung cancer transformation, epithelial-to-mesenchymal transition, and other unknown reasons (Bar & Onn, 2012, Ohashi et al., 2013, Sequist, Waltman et al., 2011, Zhang, Lee et al., 2012). To clarify those acquired resistances and better inform treatment decisions, preclinical tools to study their development are urgently needed.

Evolution-based adaptive therapy focuses on the competition between sensitive and resistant cells, using the smallest dose of drugs or even stopping the treatment (drug holiday) from maintaining its dynamic balance and achieving the purpose of prolonging survival and quality of life. Several cell line-based studies have confirmed the effectiveness of this method(Chmielecki, Foo et al., 2011). In evolution-based adaptive therapy of late-stage LUAD patients, it is still difficult to determine which patients should receive a drug holiday, and no therapeutic implications can be used for these clinical decisions.

In this study, we utilized single-cell RNA data to developed an EGFR-TKI resistance index (R-index) model with an OCLR algorithm for the first time. This model can be used to quantify the possibility of EGFR-TKI resistance. In terms of model validation, we observed that R-index could predict the dynamic changes in the number of sequential erlotinib treatment of cell lines and the outcome of osimertinib treatment xenograft mice. We also observed that R-index could predict patients’ prognosis in multiple public databases. In terms of model application, we found that the glycolysis pathway and KRAS up-regulation pathway were the predisposing factors of tumor cell resistance, and MDSC (myeloid-derived suppressor cells) is the leading cause of immunosuppression in the tumor microenvironment. And R-index can be used as a biomarker to predict the status of EGFR-TKI resistance and prognosis, providing new insights into drug resistance, and individualized treatment.

### MATERIAL AND METHODS

#### Single-cell RNA sequence (scRNA-seq) data

The scRNA-seq data(Maynard, McCoach et al., 2020) was download from Google Cloud Disk at https://drive.google.com/drive/folders/1sDzO0WOD4rnGC7QfTKwdcQTx3L36PFwX? usp=sharing. This data contains 30 advanced-stage NSCLC individual patients and 49 samples from small tissue samples as well as surgical resections. According to the medication situation, patients were divided into three states: TN (patients before initiating systemic targeted therapy, TKI naive state), RD (tumor was regressing or stable by clinical imaging, residual disease state), and PD (subsequent progressive disease as determined by clinical imaging, progression state). Smart-seq2 technology was used to extract the expression profile of single cells.

#### Cell line data

Sequential drug treatment data of PC9 cell line was obtained from Gene Expression Omnibus (GEO) database (https://www.ncbi.nlm.nih.gov/geo) under the accession number GSE149383(Aissa, Islam et al., 2021). The CCLE (Cancer Cell Line Encyclopedia) data were downloaded from the CCLE database (https://portals.broadinstitute.org/)(Barretina, Caponigro et al., 2019). The GDSC (Genomics of Drug Sensitivity in Cancer) dataset was downloaded from the GDSC database (https://www.cancerrxgene.org)(Yang, Soares et al., 2013). The CCLE and GDSC databases contain gene expression data and IC_50_ values. The IC_50_ was defined as the drug concentration for a 50% reduction of absorbance based on the survival curves.

#### Mice data

Patient-derived xenograft models of non-small cell lung cancer data were downloaded from the GEO database under the accession number GSE130160(Kita, Fukuda et al., 2019).

#### Cohort data

The OncoSG data(Chen, Yang et al., 2020) was downloaded from cbioportal at (http://www.cbioportal.org/study/summary?id=luad_OncoSG_2020). This dataset contains 305 east Asian Whole-exome and transcriptome sequencing of lung adenocarcinomas with matched normal samples. A total of 169 patient gene expression matrices were obtained, including 94 patients with EGFR mutations. The TCGA Lung adenocarcinoma data (Hoadley, Yau et al., 2018) were downloaded from cbioportal at (http://www.cbioportal.org/study/summary?id=luad_tcga_pan_can_atlas_2018). This dataset contains 510 transcriptome sequencing of lung adenocarcinomas with matched normal samples, including 54 patients with EGFR mutations. The cohort GSE31210(Okayama, Kohno et al., 2012) was downloaded from the GEO database. The dataset contains 226 Japan transcriptome sequencing data of early lung adenocarcinomas with matched normal samples, including 127 patients with EGFR mutations.

#### Resistance index (R-index) weighting matrix

We selected EGFR-mutant samples from the 49 scRNA-seq data, then performed preliminary quality control of the data, finally obtaining 14 patients, 23 samples, and 2080 cancer cells, which belonged to three different treatment time points, TN, RD, and PD. The Seurat v3(Butler, Hoffman et al., 2018) R package was used to perform the single-cell RNA-seq analysis. Cancer cells were re-clustered and visualized using a 2-dimensional t-SNE (t-distributed stochastic neighbor embedding) method. The cell lineage trajectory was inferred by Monocle2 (Qiu, Mao et al., 2017) following the tutorial. After the cell trajectories were constructed, we used the DESeq2 R package to derive DEG (Differential expressed genes) from selected branches with the p-value ≤ 0.01 and |log2FC| > 2, and got 1107 genes. A weighted 1107 genes signature array (Table S2) was yielded using one-class logistic regression (OCLR)(Sokolov, Paull et al., 2016) algorithm performed by gelnet v1.2.1 R package according to a previous study (Malta, Sokolov et al., 2018).

#### Tumor cell cluster diversity

To quantify the heterogeneity of tumor cells at three different treatment time points, we calculated the Shannon entropy of cancer cells at the corresponding time points, which captured the contribution of each tumor cell cluster(Joshi, de Massy et al., 2019). The Shannon entropy index is given by H-index=-(Σpi×log_2_(pi))/ln(N), where pi represents the relative contribution of the *i*th cluster and N is the total number of clusters. pi is obtained by dividing the count of tumor cells belonging to the *i*th cluster by the total number of tumor numbers in the treatment time point, such that Σpi=1. H-index lies between 0 (all tumor cells belonging to one cluster only) and 1 (a cluster evenly composed of all possible combinations).

#### Survival analysis

In the bulk validation cohort OncoSG, TCGA, and GSE31210, Cox proportional hazard models were used to investigate the association between R-index and patient survival. The samples were grouped into high and low expression groups by the median value. The Kaplan-Meier survival curves were plotted to show differences in survival time, and log-rank p values reported by the Cox regression models implemented in the R package survival v3.2.11 were used to determine the statistical significance.

#### fgsea analysis

Fgsea was performed using fgsea v1.10.1 R package for fast gene set enrichment analysis(Korotkevich, Sukhov et al., 2021), which had accurate standard approaches to multiple hypothesis correction allowing to make more permutations and get more fine-grained p-values.

#### ssGSEA analysis

The ssGSEA algorithm(Hänzelmann, Castelo et al., 2013) was used to quantify the relative abundance of 28 immune cell types(Charoentong, Finotello et al., 2017) and 50 hallmark gene sets(Liberzon, Birger et al., 2015) with GSVA v1.36.3 R package. The value of relative abundance was represented by an enrichment score, which was normalized to unity distribution from zero to one.

#### Cell-cell interaction analysis

We used CellPhoneDB to identify significant ligand-receptor pairs within PDB1 and RDB3. The cell-type-specific receptor-ligand interactions among cell types were identified based on the specific expression of a receptor by one cell type and a ligand by another cell type. The interaction score refers to the total mean of the individual ligand-receptor partner average expression values in the corresponding interacting pairs of cell types. The expression of any complexes output by CellPhoneDB was calculated as the sum of the expression of the component genes.

#### TIDE analysis

TIDE web application(Jiang, Gu et al., 2018) (http://tide.dfci.harvard.edu) was performed using transcriptome profiles of OncoSG, TCGA, and GSE31210 to evaluate T cell dysfunction and exclusion status.

#### Statistics

Analysis of differences between R-index median stratification groups was performed using Mann–Whitney U tests. The consistency between R-index and cell number was assessed using Spearman correlation analysis. All statistical analyses and presentations were performed using R v4.0. Statistical significance was set at p < 0.05.

## RESULTS

### 1. mRNA expression-based R-index Model

We obtained publicly multiple treatment time points Single-cell RNA sequencing (scRNA-seq) dataset(Maynard et al., 2020). Total 23 EGFR-mutant scRNA-seq LUAD samples corresponding to 14 individual patients were selected. The sample information is displayed in Table S1 and Figure S1. We derived an R-index model using one-class logistic regression (OCLR) algorithm(Sokolov et al., 2016). The OCLR-based R-index model was verified in the cell line, mice, and human cohort datasets firstly. Then, we used R-Index to explore the drug resistance mechanisms in terms of the pathway, cell interaction, and immunosuppression (Figure 1A).

**Figure 1.**
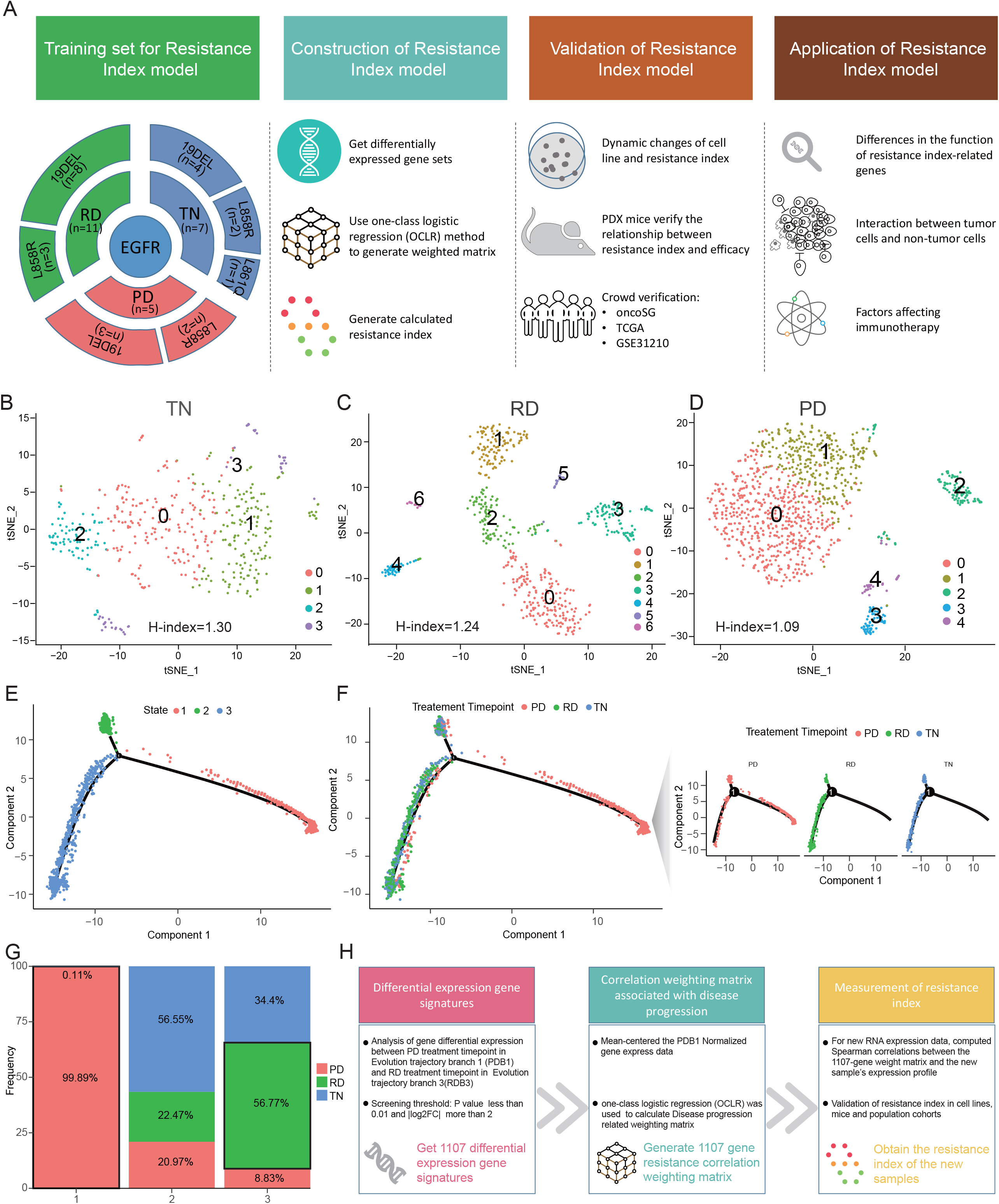
Development of the Resistance Index(R-index) (A) Overall methodology. R-index training set and its development, verification, and application. (B-D) t-stochastic neighbor embedding (t-SNE) plot and H-index of cancer cells at different treatment time points. (E) The unsupervised transcriptional trajectory of cancer cells from Monocle (version 2), colored by cell states and (F) treatment timepoint. (G) The relative proportion of cancer cells for three treatment timepoints in each state as shown in (E). (H) The workflow for the development and application of the R-index model.

### 2. Identification of R-index signature genes reflecting the drug resistance transcriptional heterogeneity of cancer cells

A total of 2080 cancer cells were retained after quality control filtering. All cancer cells at different treatment time points were re-clustered and visualized using the t-distributed stochastic neighbor embedding (t-SNE) method, and H-indexes were also calculated respectively. The cell clusters at TN, RD and PD were 4, 7, and 5, respectively, which indicates cancer cells heterogeneous in different treatment time points (Figure 1B-1D). To quantify the heterogeneity among cancer cells, we calculated the diversity index of each cluster. The results showed that the H-index order was TN>RD>PD. We proposed a hypothesis that drug intervention might affect tumor cell differentiation due to environmental screening, making drug-resistant cells obtain more competitive advantages than sensitive cells, reducing the diversity of cancer cell composition.

Trajectory analysis was performed with monocle software to project all cancer cells to explore the heterogeneity and the cells that play a major role in governing the tumor progression (Figure 1E). Indeed, transcriptional states in the trajectory revealed differentiation of cancer cells at different treatment time points (Figure 1F). Firstly, cancer cells were located in separate trajectory branches, which marked their distinct differentiation states. Secondly, branch 1 was mainly occupied by PD timepoint cells, branch 2 by TN timepoint cells, and branch 3 by RD timepoint cells (Figure 1G). Lastly, the differentiation direction of PD timepoint cells was different from TN and RD sample cells.

To identify transcriptional signatures defining cellular resistance status in the trajectory, we compared differentially expressed genes specific to PD in branch 1 (PDB1) and RD in branch 3 (RDB3). 1107 candidate genes were filtered out, in which 348 genes were upregulated in PDB1 and 759 genes were upregulated in RDB3. Considering that these genes may contain biological features that potentially distinguished the drug-resistant states of cells, we applied one-class logistic regression (OCLR) algorithm to build a model which was trained on PDB1 cells and produced a weighted 1107 gene matrix to extract transcriptomic features of the drug resistance signature. The R-index score was defined as the Spearman correlation coefficient of the 1107 gene signature matrix and a new validation data set (Figure 1H).

### 3. Assess the predictive ability of R-index for resistance status in cell lines and mice

Given that the R-index is hypothesized to quantify the degree of resistance of samples to EGFR inhibitors, we examined the R-index dynamic changes of PC9 cells treated with erlotinib(Aissa et al., 2021). The data were obtained from PC9 cells subjected to chronological erlotinib treatment for 0, 1, 2, 4, 9, and 11 days. Then drop-seq technology was applied to evaluate single-cell gene expression profiles. PC9 cells contained the exon 19 deletion in the EGFR gene and can be used to simulate patient-related intrinsic and acquired TKI (Tyrosine Kinase Inhibitor) resistance in vitro(Sharma, Lee et al., 2010). Erlotinib, an irreversible first-generation EGFR-TKI, exerts cytostatic and cytotoxic effects on PC9 at 2 μM, which is used as first-line treatment for patients with EGFR mutation-positive(Planchard, Popat et al., 2018). The results showed that the R-index value changed significantly with the extension of the medication time but nonlinearly (Figure 2A). To test if similar patterns of change exist in cancer cell populations and R-index, we used a line graph to illustrate the dynamic changes of the number of cells and the R-index value and found that they had opposite trends (Figure 2B) with a strong negative correlation (Spearman correlation r = -0.79, Figure 2C).

**Figure 2.**
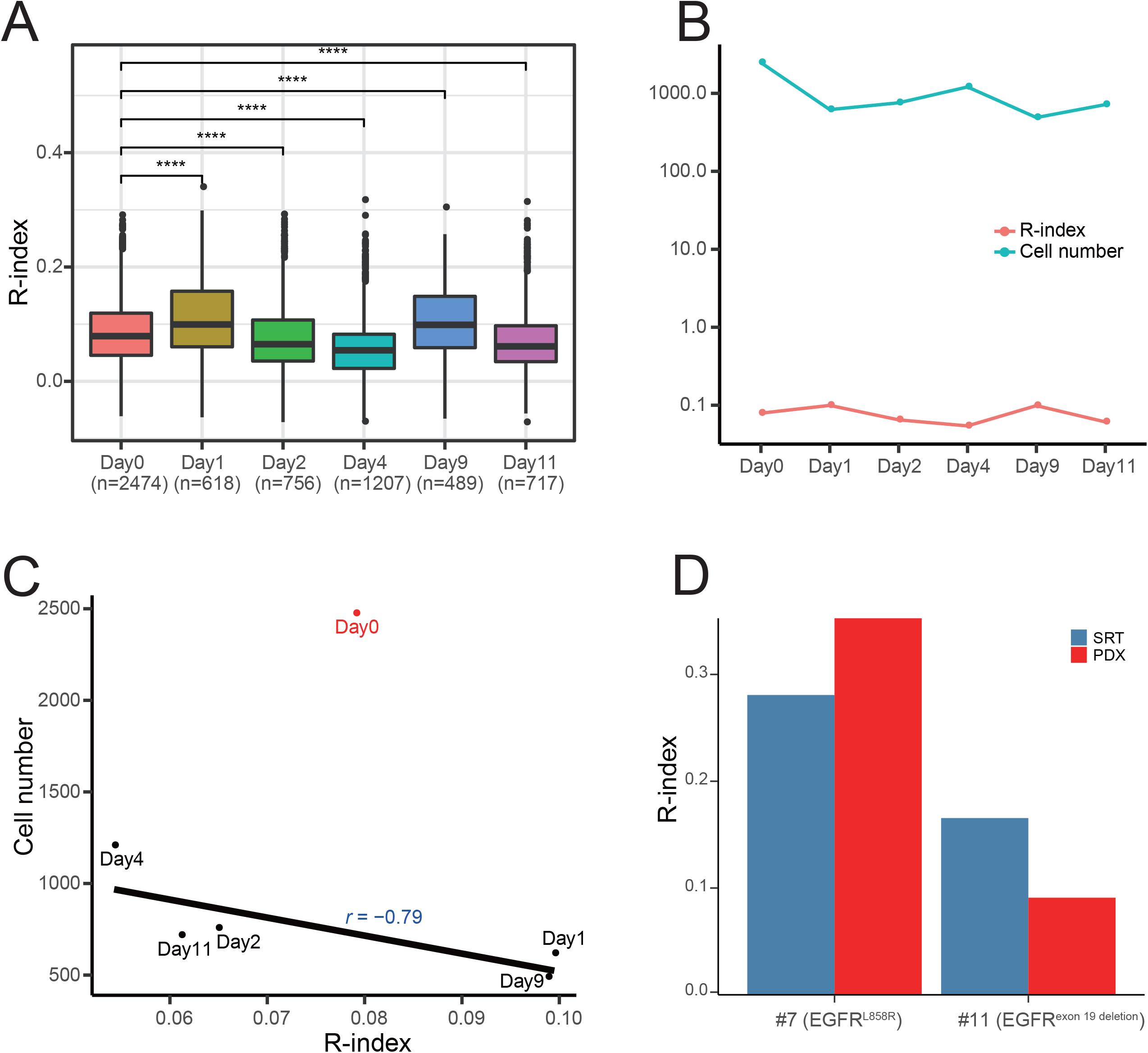
Validation of R-index in cell lines and mice. (A) R-index boxplot for the set of consecutive cell lines for single-cell RNA-seq. Day 0 are untreated cells, and Day 1 through Day 11 is the duration of treatment with 2 mM erlotinib. (B) Simultaneous display of R-index and cell number changes line graph and (C) Correlation diagram. (D) R-index changes in mice between treatment-naive and osimertinib treatment in vivo. SRT, surgically resected tumors; PDX, patient-derived xenografts.

Since the cells in culture lacked many in vivo interactions, the R-index was applied to investigate the response of mice to EGFR TKI with data from a xenograft study(Kita et al., 2019). Patient-derived xenograft (PDX) models were built by implanted small pieces (3-5 mm) of adenocarcinomas specimens from patients’ surgically resected tumors (SRT) with EGFR-activating mutations (#7, #11) into the subcutaneous flank tissue of female SHO mice (Crlj: SHO-PrkdcscidHrhr, Charles River). Tumor size was measured with calipers once a week, and the mice were treated by oral gavage with 25 mg/kg per day of osimertinib when tumor volume exceeded 500 mm^3^. When tumor volume reached 1500 mm^3^, mice were killed, and tumors were implanted into new mice. Tumor fragments #7 had EGFR L858R mutation, and #11 had EGFR exon 19 deletion mutation. The R-index was calculated based on gene expression (Figure 2D). As PDX tumor in case #7 regrew during the continuous osimertinib treatment at the fifth passage, we calculated and found that its R-index was higher than that of SRT. The PDX tumor in case #11 was cured at the third passage, and as expected, the R-index had an opposite result compared with case #7.

### 4. Correlations of R-index with overall survival in the external cohort data

After validating the predictive power of the R-index in the cell line and mice, we further proved that the R-index could also quantify primary patients’ resistance status and predict prognosis in multiple datasets. We first calculated the R-index of EGFR-mutant samples from the OncoSG database and dichotomized patients into two equal-size groups using the median R-index as the threshold. We found that the high R-index group showed a significantly shorter overall survival time than the low R-index group (p = 0.008, Fig 3A). Similarly, in this total cohort, a high R-index group was also associated with worse outcomes (p=0.001, Fig 3B). To further evaluate the results, another two lung adenocarcinoma gene expression datasets were examined. The median resistance index was still used as the stratification threshold. As expected, in the datasets of the TCGA cohort and GSE31210 cohort, we also observed that the high R-index group showed a shorter overall survival time than the low R-index group, regardless of whether it is in EGFR-mutant patients ( p=0.07, TCGA, Figure 3C; p=0.07, GSE31210, Figure 3E) or the entire cohort (p<0.001, TCGA, Fig 3D; p<0.001, GSE31210, Fig 3F).

**Figure 3.**
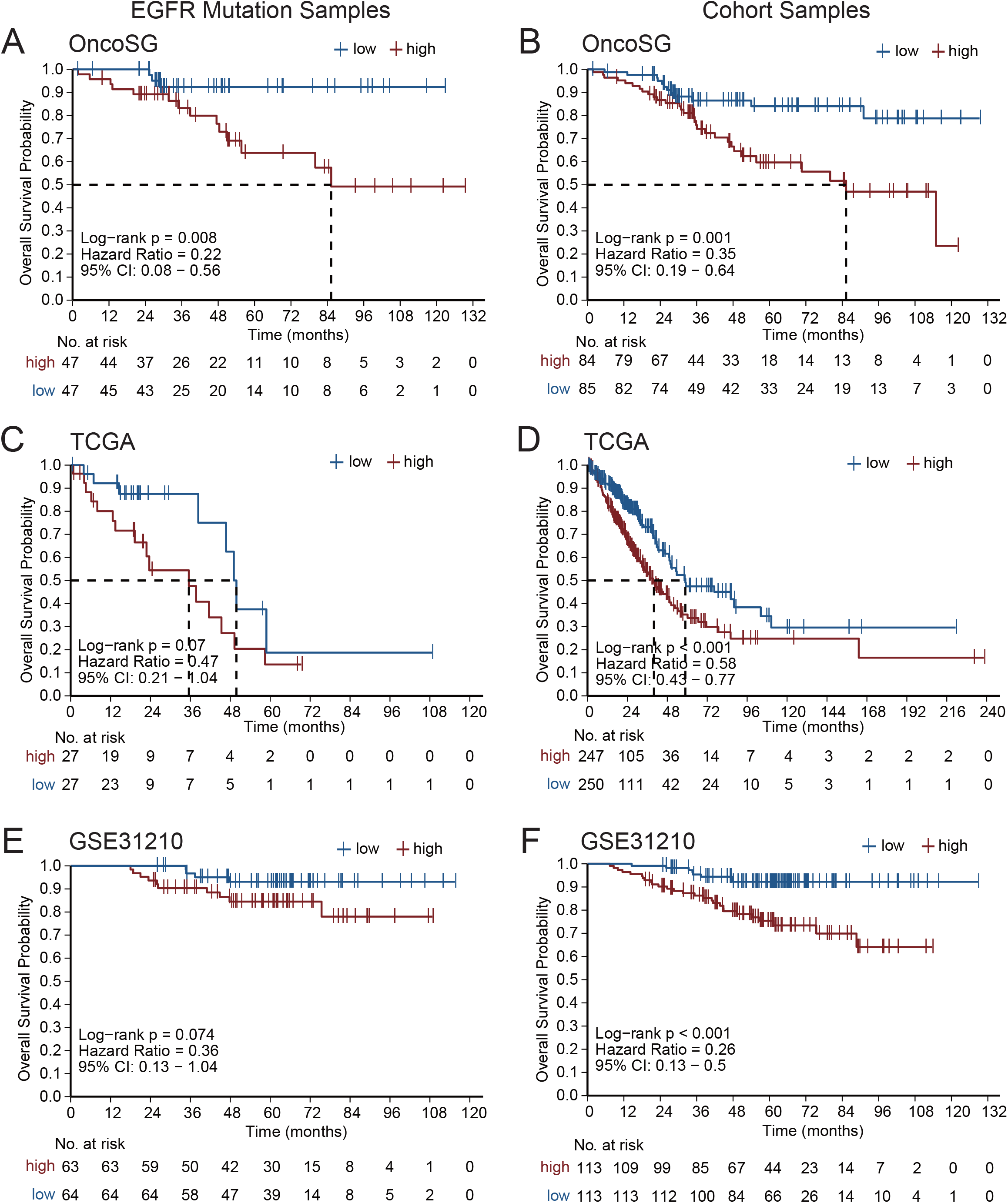
Validation of R-index in the human cohort. EGFR mutation samples and the entire cohort samples in the OncoSG (A-B), TCGA_LUAD (C-D), and GSE31210 (E-F) databases to verify the relationship between R-index and prognosis, and the threshold is the median R-index of the corresponding cohort.

### 5. Analysis of R-index signature genes functional features in hallmark gene set

Our analyses demonstrated that the R-index was associated with the resistance status of cell lines and mice. And the prognosis of NSCLC patients can also be evaluated according to R-index stratified analysis. Therefore, we hypothesized that the substantial differences in the tumor expression pattern between PDB1 and RDB3 could be caused by differences in cancer cells. To explore these differences, a fgsea script was performed to analyze R-index signature genes using the hallmark gene set in MSigDB v7.4. The analysis yielded 9 significantly enriched gene sets. Metabolism-related glycolysis and signaling-related KRAS signaling up gene sets were significantly positive-enriched in PDB1 (Figure 4A), and the volcano plot highlighted related differential genes (Figure 4B).

**Figure 4.**
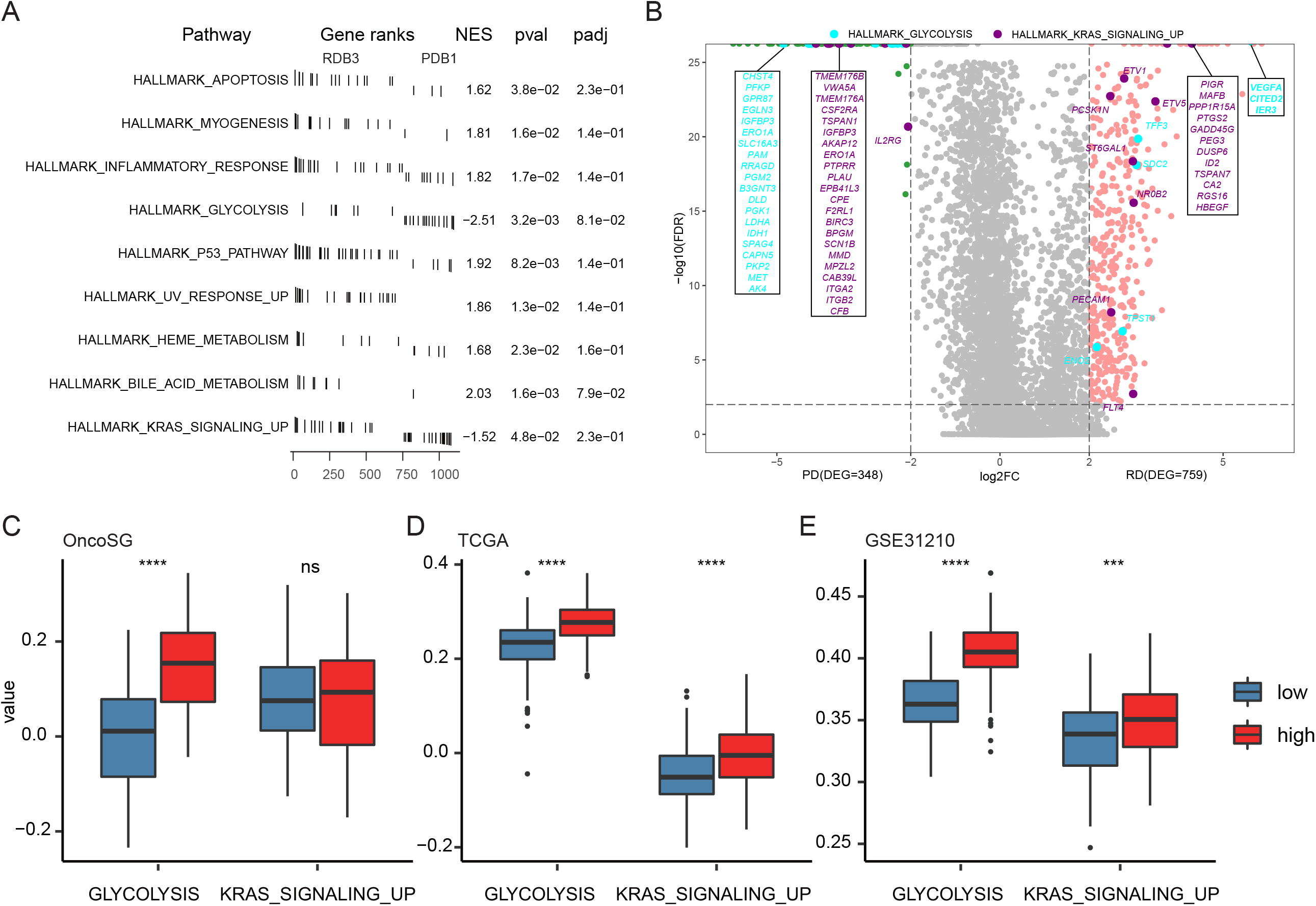
Enrichment of R-index-related functional gene sets. (A) R-index related function enrichment with hallmark gene sets from MSigDB analyzed by fgsea. (B) Volcano map to show the detailed gene sets of glycolysis pathway and KRAS up-regulation pathway. (C) Glycolysis and KRAS up-regulation gene sets are orthogonally verified in public databases.

In contrast, pathway-related apoptosis, immune-related inflammatory response, and proliferation-related p53 pathway were significantly positive-enriched in RDB3 (Figure 4A). To verify whether there were consistent results in the external database, we estimated the value of each hallmark’s ssGSEA profile in OncoSG, TCGA, and GSE31210. In line with expectations, glycolysis and KRAS_signaling_up expressed significantly higher in the high R-index group using median stratification (Figure 4C-E). In addition, the expression of the epithelial-mesenchymal transition (EMT) gene set was also higher in the high R-index group (Figure S3).

### 6. Intercellular communication networks analysis

To investigate potential interactions between cancer cells and other immune cells in the tumor microenvironment (TME), we performed cell-cell communication analysis using CellPhoneDB [38], a Python program calculating the interaction between the receptors and ligands. Based on research purposes, we divided tumor cells into three types according to their evolutionary trajectories, namely PDB1 cells and RDB3 cells used in the previous analysis and Other_cancer_cells that did not contain these two types of cells. Enriched receptor-ligand interactions network diagrams were derived based on the expression of receptors and the corresponding ligand between two connected cell types for demonstrating their extensive communication (Figure 5A). To further investigate the interactions that occur in the TME, we utilized receptor-ligand pairs to calculate the strengths of the interactions. Cancer cells showed close interactions with fibroblast, MF-Monocytes, endothelial, and dendritic cells in both PDB1 and RDB3. When using the odds ratio to normalize the receptor-ligand pairs of the two time points, we found that the Neutrophils, B-cells-M (B memory cell), B-cells-PB (B plasma cell), and T-cells had a higher ratio in the PDB1 time point (Figure 5B).

**Figure 5.**
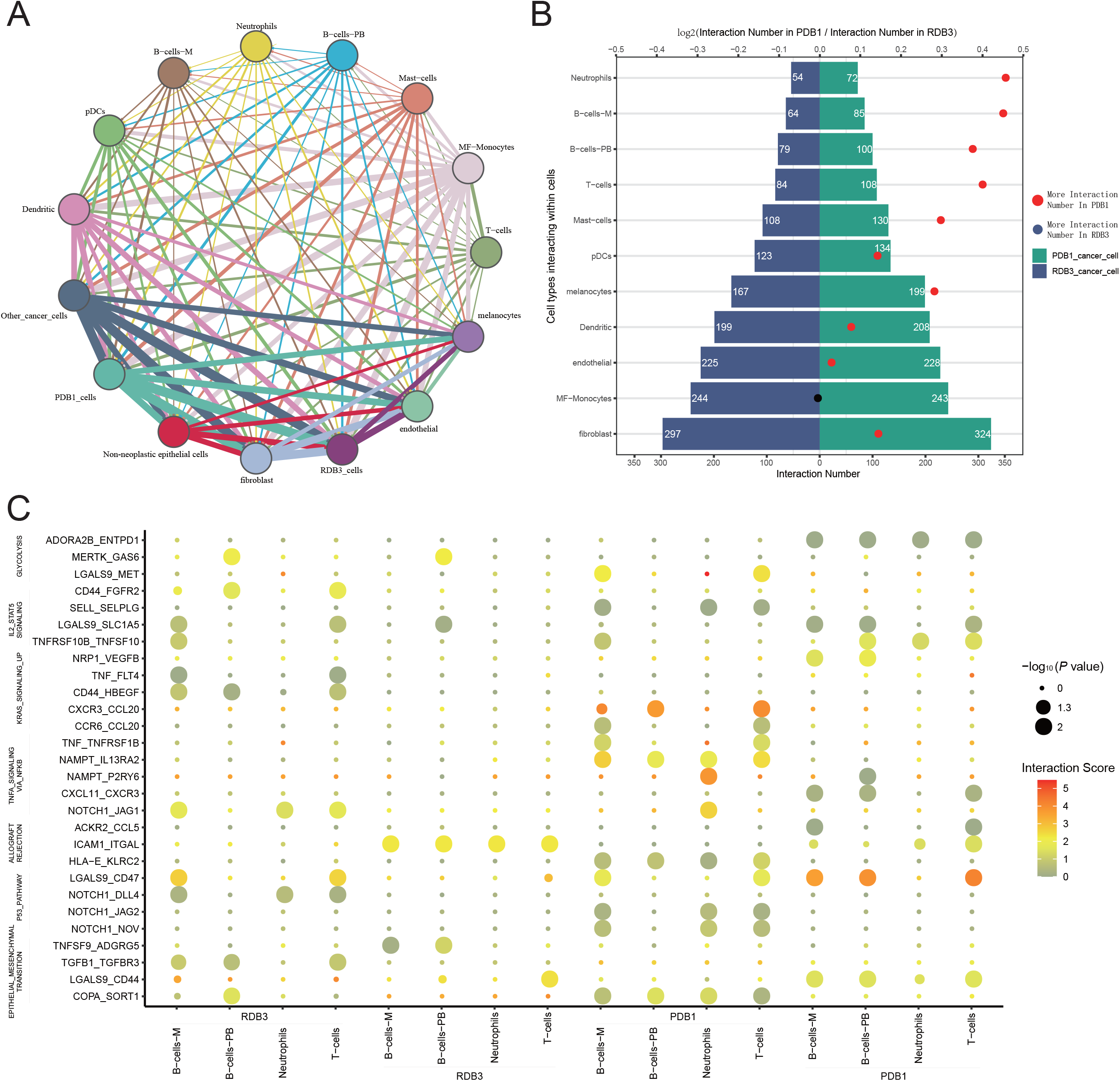
Interaction between cancer cells and tumor microenvironment cells. (A) Intercellular communication between cancer cells and others cells. Each line’s color and thickness indicate the ligands and receptors’ connection and proportion, respectively. (B) Bar chart showing the number of ligand-receptor pairs in cancer cells and other cells in PDB1 and RDB3 samples. and the dots represent their ratio, the ratio above 1 in red and below 1 in black. (C) Overview of selected ligand-receptor interactions in hallmark gene set between cancer cells and top-four ratio cell type.

Based on the results of quantitative analysis of the Receptor-ligand pair, we used bubble diagrams to specifically show the interactions between two timepoint tumor cells (PDB1 and RDB3) and four types of immune-related cells (Neutrophils, B-cells-M, B-cells-PB, and T-cells) (Figure 5C). And finding that immunosuppressive-related receptor-ligand gene ADORA2B(Liu, Kuang et al., 2020), ENTPD1(Moesta, Li et al., 2020), CXCR3(Chow, Ozga et al., 2019), LGALS9(He, Jia et al., 2019) showed strong regulatory relationships with PDB1.

### 7. Suppressive immune microenvironment primed by myeloid cells

Finally, we explored the treatment roles of immune signatures. First, immune surveillance and escape signatures were used to quantify the features of immune infiltration(Sun, Wu et al., 2021) between RDB3 and PDB1, and it was observed that PDB1 had higher immune escape ability (Figure 6A). Second, we examined the expression of immune checkpoint inhibitor-related genes CD274 and CTLA4 in the public database, observing that the high R-index group had a significantly higher expression level with median stratification (Figure 6B). In addition, the TMB status had consistent results (Figure S4). Third, considering that the PD had immunosuppressive features(Maynard et al., 2020), we used the TIDE (Tumor Immune Dysfunction and Exclusion) algorithm to identify factors that excluded T cell infiltration into tumors from the large patient cohort. As shown in Figure 6, Among three cell types related to T cell exclusion, there was no significant difference between TAM.M2 (the M2 subtype of tumor-associated macrophages) and CAFs (cancer-associated fibroblasts) when using R-index median stratification. However, MDSC was significantly higher in the high R-index group (Figure S5). We also observed that the ROS (reactive oxygen species) pathway (Figure S3) was significantly higher in the high R-index group.

**Figure 6.**
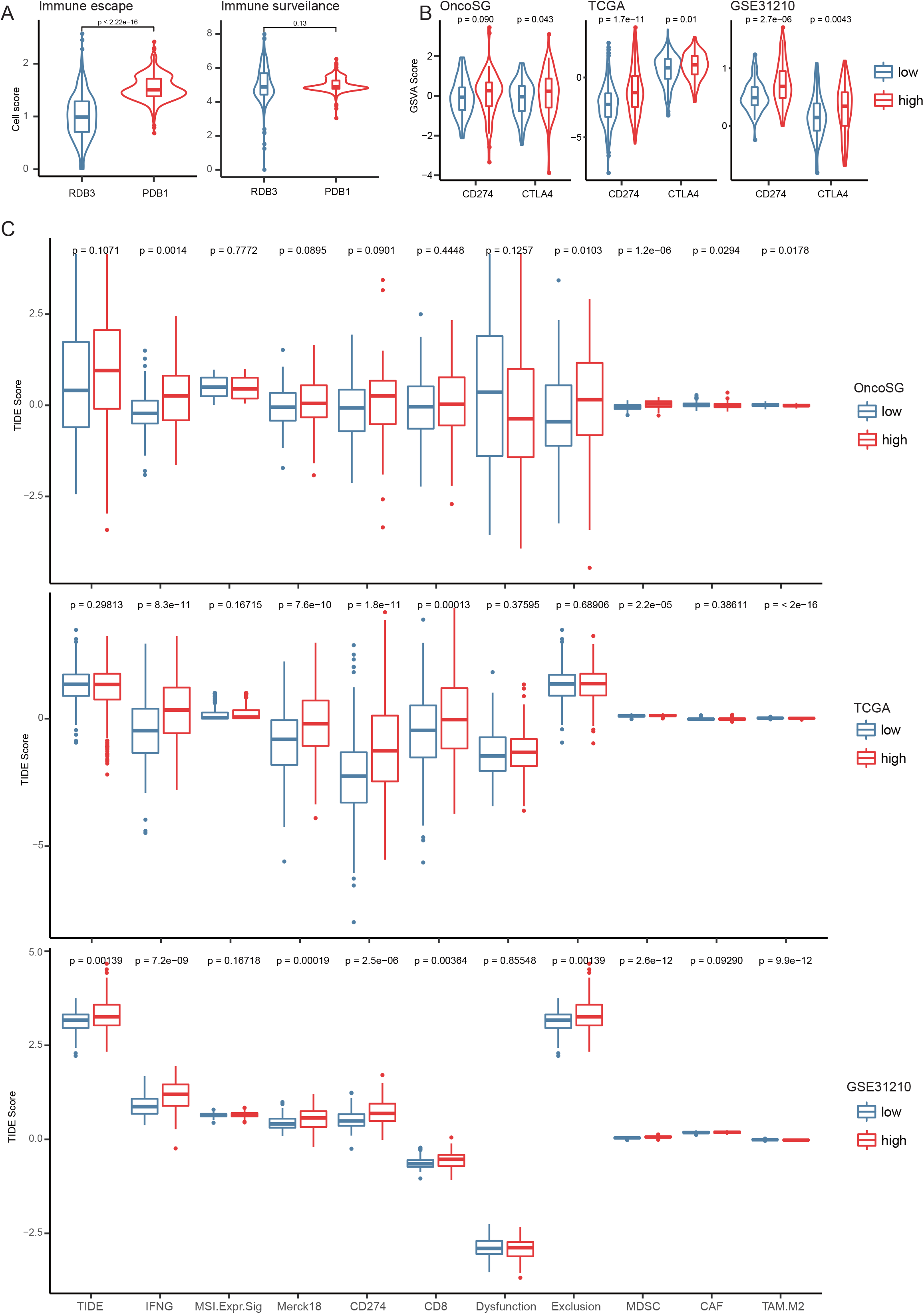
The contributions of R-index stratification to the immune escape. (A) Comparison of Immune escape and Immune surveillance signature scores of PDB1 and RDB3. (B) Comparison of *CD274* and *CTLA4* gene expression. (C) TIDE score in external databases based on R-index stratification.

## DISCUSSION

EGFR-TKIs dramatically increased the five-year survival rate of late-stage patients harboring EGFR-activating mutations. However, patients acquired resistance inevitably after a period of target treatment. Since drug resistance leads to the heterogeneity of tumors, the status of EGFR-TKI resistance is complex, and the mechanism is varied. At present, the exploration of drug resistance mechanisms mainly focuses on off-target alterations or overexpression of several genes. The study in RNA expression level has not been well explored before. An effective RNA-based biomarker for predicting EGFR-TKI resistance is helpful for medical care. Using the single-cell RNA-seq data at three different treatment time points obtained from patients, we developed the R-index model trained from EGFR-targeted therapies samples at the single-cell level to quantify the possibility of resistance. Then the model was validated in vitro and in vivo for predicting resistance. For the proportion of resistant cells in the primary patients that may affect the prognosis due to the mediation of resistance, we further validated the model in public databases with EGFR-mutant patients and the entire cohort. Finally, we explored the value of the R-index in practical applications from the three dimensions of tumor cell expression pattern, cell interactions, and immune responses.

In an in vitro cell line verification scenario, we adopted PC9 cells that received intermittent erlotinib treatment. After calculating the number of cells and the R-index of each cell on the 0, 1, 2, 4, 9, and 11th days, we found that neither the number of cells nor the average R-index changed linearly. However, there was a clear negative correlation between them. In general, the number of cells could reflect their own growth and proliferation states to some extent. Under the selective pressure of the drug, drug-sensitive cells were eliminated. Then drug-resistant tumor cells remained after treatment proliferates rapidly due to the expansion of living space. Evolution-based treatment(Zhang, Cunningham et al., 2017) or the “drug holiday” phenomenon in treatment(Song, Yu et al., 2014) as a conceptual treatment strategy has developed for many years(Aktipis, Kwan et al., 2011) based on Darwinian dynamics of intratumoral heterogeneity. The content of this theoretical framework is that cancer cells are commonly heterogeneous and contain both resistant cells and sensitive cells. Environmental selection forces (E.g., medication) can establish a new balance in line with Darwin’s theory(Gatenby & Brown, 2020). During the drug holiday, without drug interference, drug-sensitive tumor cells could quickly proliferate and compete with the drug-resistant tumor cells (Gatenby, Cunningham et al., 2014, Shen, Chu et al., 2008). In a previous study, passaging drug-resistant cells displayed slower growth kinetics compared to drug-sensitive parental cells and could restore drug sensitivity after drug withdrawal(Chmielecki et al., 2011). Clinically, many reports about salvage treatment demonstrated that patients who had acquired resistance before could re-respond to EGFR-TKI re-challenge after the drug holiday (Oh, Ban et al., 2012, Watanabe, Tanaka et al., 2011, Yamaguchi et al., 2019). When the drug was applied after the drug holiday, the number of drug-sensitive cells reversed and EGFR-TKI tolerant patients regain clinical benefit, which meant the drug resistance states of tumor cells changed dynamically in the process of alternate selective pressure.

Traditional approaches to cancer therapy commonly produced a partial or complete response but were inevitably followed by disease progression. Then drugs of subsequent treatment were tried. However, the outcome was always disappointing because resistant cells had taken an absolute advantage. Evolutionary dynamics treatment strategies attempted to maintain the possibility of long-term tumor control by slowing the proliferation of the resistant population. For this purpose, minimum dose administration combined with periodic dosing and withdrawal was performed to maximize the dynamic balance of resistant and sensitive cells and maintain tumor homeostasis. Therefore, to assist in making the clinical decision, a clear biomarker is urgently needed to monitor the dynamic changes of the resistance status of tumor cells. Here we for the first time proposed an RNA-based index, R-index, as a quantification biomarker of EGFR-TKI resistance status to support evolutionary dynamics treatment in EGFR-mutant patients. And our method of designing the R-index may be utilized in other types of tumors in the future.

In the second part of our research, we applied the R-index to CCLE (Cancer Cell Line Encyclopedia) EGFR mutation(Barretina, Caponigro et al., 2012), and GDSC (Genomics of Drug Sensitivity in Cancer) (Yang et al., 2013) cell lines databases. We observed an interesting phenomenon: the IC_50_ value of the high R-index group was significantly lower than that of the low R-index group (groups were divided by the median R-index) (Figure S2A-B). This result could be explained by the slower growth of the drug-resistant cell, leading to fewer fractions under mild conditions. A study showed that T790M-containing resistant cells grew slower than parental cells(Chmielecki et al., 2011). We used the proliferation score to evaluate the proliferation status of the cells of day 0 and found that the score of the high R-index group was significantly lower than that of the low R-index group (Figure S2C).To assess the relative abundance of resistant cells, we used an empirical threshold of 0.26 from the R-index of CCLE (median value 0.254) and GDSC (median value 0.262) to divide the resistant cell in the cell population Day 0. As expected, the proportion of cells greater than the threshold was less than 5% (Figure S2D).

When we further used in vivo PDX model to evaluate the response of EGFR-TKI. Two EGFR mutations (L858R or exon 19 deletions) Patient-derived tumors were transplanted into mice and continuous oral treatment with Osimertinib. We calculated the R-index of PDX mice and control SRT tissue samples, respectively, and observed that R-index increased in resistant mice and decreased in cured mice, which confirm again that the R-index is suitable for the prediction of clinical responses to targeted drugs and maybe as an evaluation tool for the efficacy of novel treatment.

Tumor heterogeneity is a pathological property where tumors and their surrounding microenvironment were different among patients, determining the difference in treatment. Patients management during the treatment process was also a challenge. To corroborate the R-index that can be used as a prognostic biomarker, we compared the clinical outcomes of patients with EGFR mutation and the entire cohort in the external database using stratification of the median R-index. The prognosis of the high R-index group was worse than that of the low R-index group, which was consistent with prior results. Because of the biological plasticity of tumor cells, early effective treatment often produces drug resistance in the later stage(Yuan, Norgard et al., 2019). Several studies have explored the mechanisms by comparing baseline and re-biopsy tissue specimens(Kobayashi, Boggon et al., 2005, Yu, Arcila et al., 2013). However, performing surgery or biopsy to obtain tissues from relapsed patients has many limitations, even in large, well-designed clinical trials(Fukuoka, Wu et al., 2011, Lee, Park et al., 2014). Identifying the molecular mechanism of acquired resistance and developing relevant drugs are needed for effective posterior treatments. We consider that the R-index might be able to indirectly explore the resistance mechanism of large population cohorts by quantifying and comparing the resistance of primary patients.

We tried to prove the above hypothesis through 3 aspects. First of all, we found that the R-index built on the weighted 1107 gene signatures showed satisfactory performance of distinguishing tumors with resistance expression patterns in both laboratory control conditions and external cohort datasets. Fgsea analysis showed that the glycolysis metabolism and KRAS upregulate pathway was significantly enriched in PDB1. The Warburg effect describes the phenomenon that tumor cells increased utilization of glycolysis rather than oxidative phosphorylation to dominates ATP production despite adequate physiological oxygen conditions(Warburg, 1956). On the one hand, glycolysis can depress tumor cell differentiation and apoptosis to promote proliferation(Tomiyama, Serizawa et al., 2006, Vander Heiden, Cantley et al., 2009). On the other hand, glycolysis produces excessive lactate to creates an acidic tumor microenvironment that promotes invasion and migration(Hirschhaeuser, Sattler et al., 2011). Inhibition of increased lactic acid production can seriously affect disease progression (Xie, Hanai et al., 2014). Several studies demonstrated that increased glucose metabolism in tumor cells is associated with resistance to EGFR-TKI treatment. The combined use of glucose metabolism inhibitors could be an effective therapeutic strategy for patients with higher R-index (Kim, Yun et al., 2013, Suzuki, Okada et al., 2018, Tamada, Nagano et al., 2012). The oncogene *RAS* was first revealed through its ability to promote glycolysis and resistance to targeted therapy(Kitajima, Asahina et al., 2018, Racker, Resnick et al., 1985). As the downstream mechanism of the EGFR signaling pathway, the activation of the KRAS-RAF-ERK pathway plays an important role in the malignant transformation of normal cells. KRAS and its downstream stand-out signaling pathways, such as MAPK, PI3K, and RAL-GDS, has been used as important sources to discover treatment opportunities(Kitajima et al., 2018, Lito, Rosen et al., 2013).

In the third major part of our study, we analyzed the interaction of receptors and ligands between tumor cells and microenvironment cells from quantitative and qualitative aspects. In terms of quantity, we found that Neutrophils, B-cells-M, B-cells-PB, and T-cells interacted more closely with PDB1. The neutrophil is an essential component of regulating adaptive immune responses by expressing a vast repertoire of cytokines(Scapini & Cassatella, 2014). In most human tumors, tumor-associated neutrophils infiltration was associated with poor prognosis(Shaul & Fridlender, 2019). Another type of neutrophils related to immunosuppressive activity differentiates into MDSC, a heterogeneous population of mostly immature myeloid cells(Bronte, Brandau et al., 2016). In the quantitative analysis, we also detected the receptor and ligand pairs related to immunosuppression, like ADORA2B-ENTPD(Chen, Akdemir et al., 2020). In addition, the single-cell data also confirmed that the PD treatment time point contained an immunosuppressive tumor microenvironment(Maynard et al., 2020).

Finally, to refine the relationship between T cells and tumor cells, we first used the immune escape score [43] to compare the differences between PDB1 and RDB3 macroscopically. The results showed that the PDB1 had a higher immune escape ability. We then compared tumor cell immune checkpoint gene expression levels between patients with high or low R-index in public databases. Consistent with former results, the expression of PDL1 and CTLA4 genes in the median stratified high R-index group was significantly higher than that in the low R-index group, and the TMB had the same trend. Several EGFR-TKI studies reported that the PD-L1 expression and TMB increased in resistant samples (Isomoto, Haratani et al., 2020, Peng, Wang et al., 2019), indicating the R-index had a prediction value in EGFR-TKI treatment.

With the TIDE algorithm, we distinguish whether the immune escape was caused by T cell dysfunction or exclusion on the public database. And it was indicated that MDSC-mediated immune exclusion might be a factor of immune escape. The supporting data came from the above-mentioned cell interactions and MDSC score and the IN-gamma (INFG) score, and ROS pathway scores were significantly higher in the high R-index group. As we know, MDSC also played an important role in the resistance of tumor cells against the immune system to specific therapies(Domagala, Laplagne et al., 2021, Li, Salehi-Rad et al., 2021, Weber, Fleming et al., 2018).

There are still several limitations in our study. First, although the single-cell data provide a high resolution to study the cell characters of the tumor, the sample size involved is limited. Due to population heterogeneity and the potential of the dropout phenomenon, it may contribute to biased results during the calculation of the weighting matrix. Another shortcoming is that the bulk RNA data of EGFR-TKI resistance tissues were scarce, which precluded us from setting a cohort to analyze the effectiveness of the R-index of our findings. A sufficient number of patients will be enrolled, and EGFR-TKI treatment clinical outcomes will be collected in the follow-up study, and these limitations will be overcome; therefore, deeper mechanism mining and clinical verification will be realized.

## DATA AVAILABILITY

The scRNA-seq data was download from Google Cloud Disk at https://drive.google.com/drive/folders/1sDzO0WOD4rnGC7QfTKwdcQTx3L36PFwX? usp=sharing

Sequential drug treatment data of PC9 cell line was obtained from Gene Expression Omnibus (GEO) database (https://www.ncbi.nlm.nih.gov/geo) under the accession number GSE149383

The CCLE (Cancer Cell Line Encyclopedia) data were downloaded from the CCLE database (https://portals.broadinstitute.org/)

The GDSC (Genomics of Drug Sensitivity in Cancer) dataset was downloaded from the GDSC database (https://www.cancerrxgene.org)

Patient-derived xenograft models of non-small cell lung cancer data were downloaded from the GEO database under the accession number GSE130160

The OncoSG data was downloaded from cbioportal at (http://www.cbioportal.org/study/summary?id=luad_OncoSG_2020)

The TCGA Lung adenocarcinoma data were downloaded from cbioportal at (http://www.cbioportal.org/study/summary?id=luad_tcga_pan_can_atlas_2018)

The validation cohort data were obtained from GEO under the accession number GSE31210

Software:

Seurat https://github.com/satijalab/seurat Monocle2

http://bioconductor.org/packages/release/bioc/html/monocle.html

gelnet (v1.2.1)

https://cran.r-project.org/web/packages/gelnet/index.html

R 4.0.5

https://www.R-project.org fgsea

(v1.14.0)

http://bioconductor.org/packages/release/bioc/html/fgsea.html DESeq2 http://bioconductor.org/packages/release/bioc/html/DESeq2.html

Survival

https://github.com/therneau/survival

ggplot2

https://cran.r-project.org/web/packages/ggplot2/index.html

GSVA

http://bioconductor.riken.jp/packages/3.0/bioc/html/GSVA.html

CellPhoneDB

https://github.com/Teichlab/cellphonedb

TIDE

http://tide.dfci.harvard.edu

## FIGURES LEGENDS

**Figure S1.**
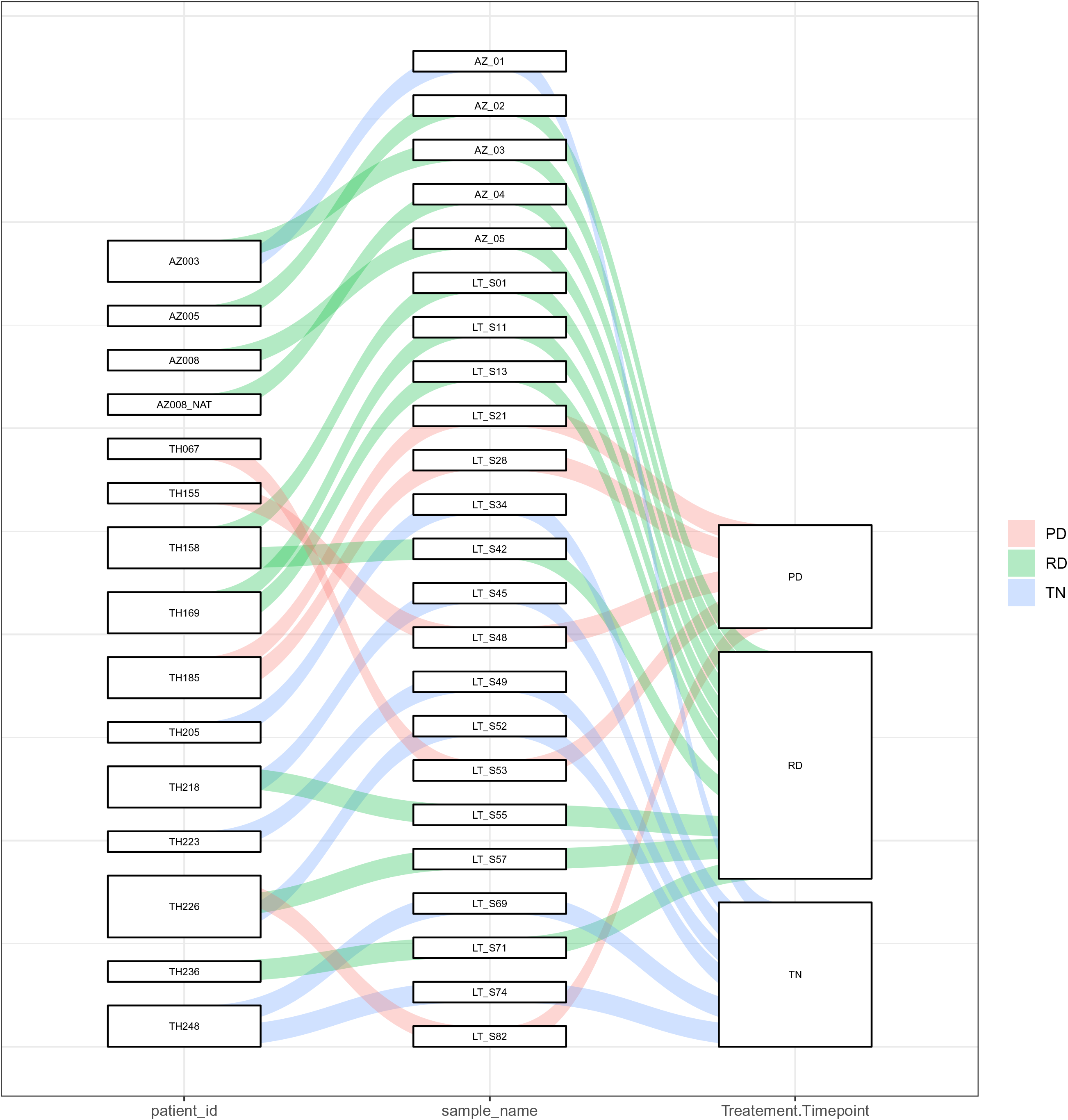
Sankey diagram of patients, samples, and treatment time points.

**Figure S2.**
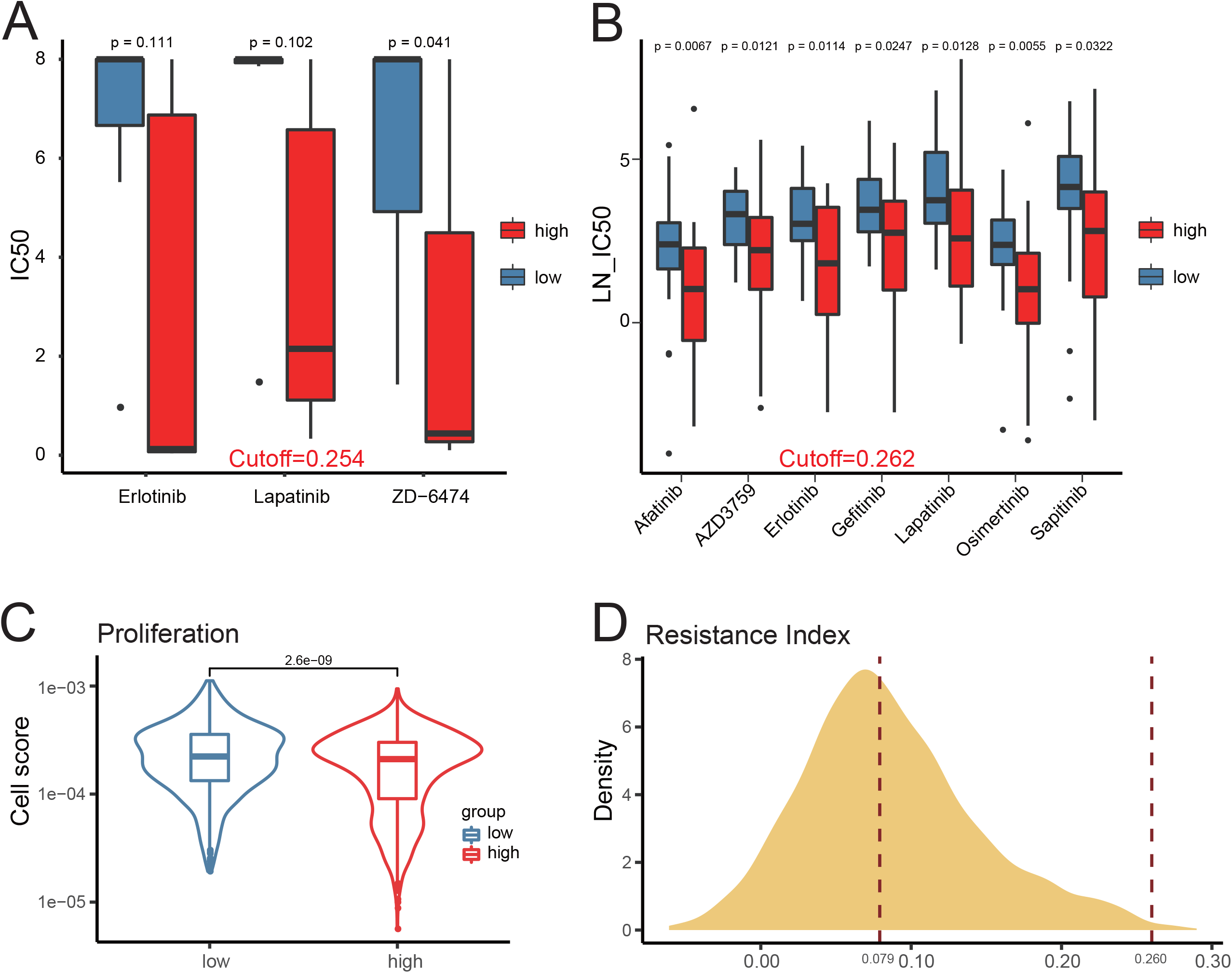
R-index stratified analysis in cell lines database. (A) Comparing IC50 difference between CCLE EGFR-mutant cell line and (B) GDSC cell line according to the median R-index stratification. (C) Violin Chart of cell proliferation signature score for the day 0 cell line in Figure 2A according to the median R-index stratification. (D) The R-index density distribution chart for the day 0 cell line in Figure 2A, and the dotted lines represent the median and the empirical threshold respectively.

**Figure S3.**
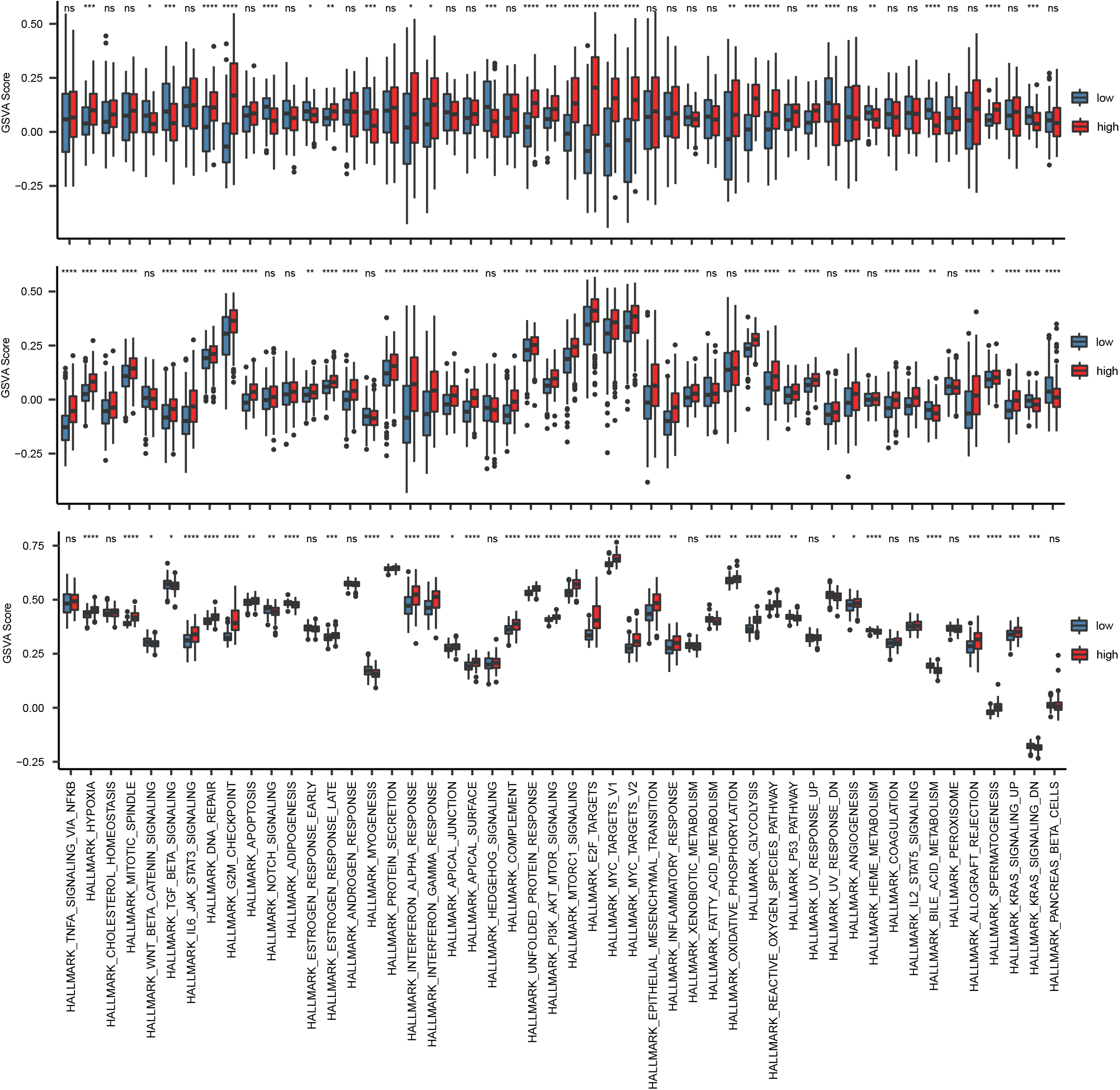
Stratified analysis of the Hallmark gene set in the external database. Comparative analysis of hallmark gene set ssGSEA score according to the median R-index stratification in OncoSG, TCGA_LUAD, and GSE31210 database.

**Figure S4.**
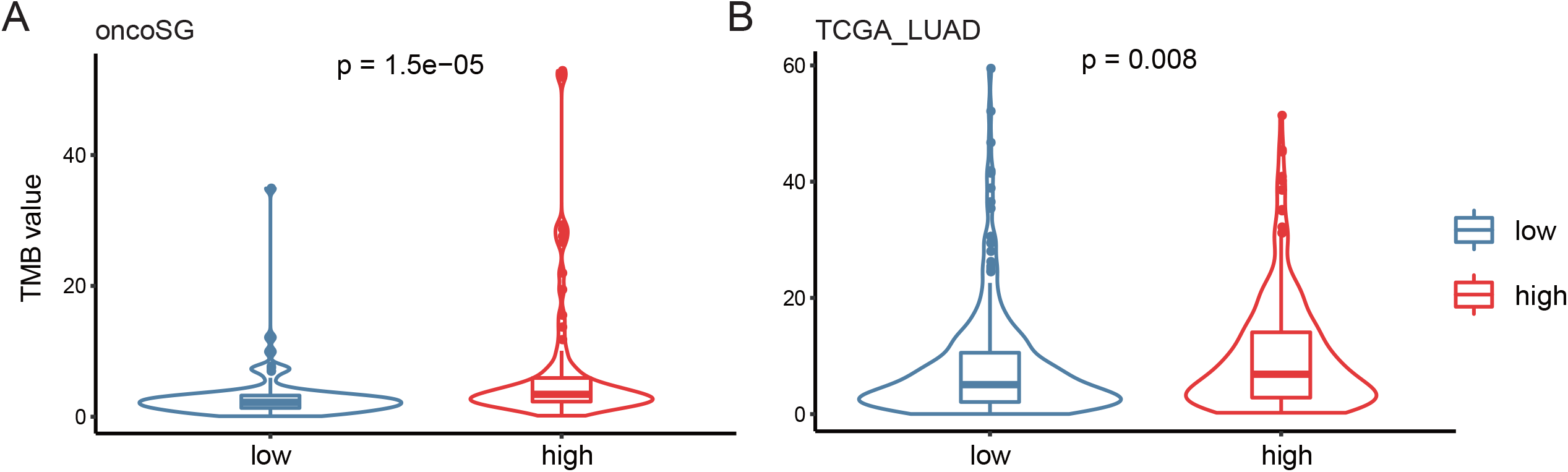
TMB features. Comparative analysis of TMB value according to the median R-index stratification in the OncoSG and TCGA_LUAD database.

**Figure S5.**
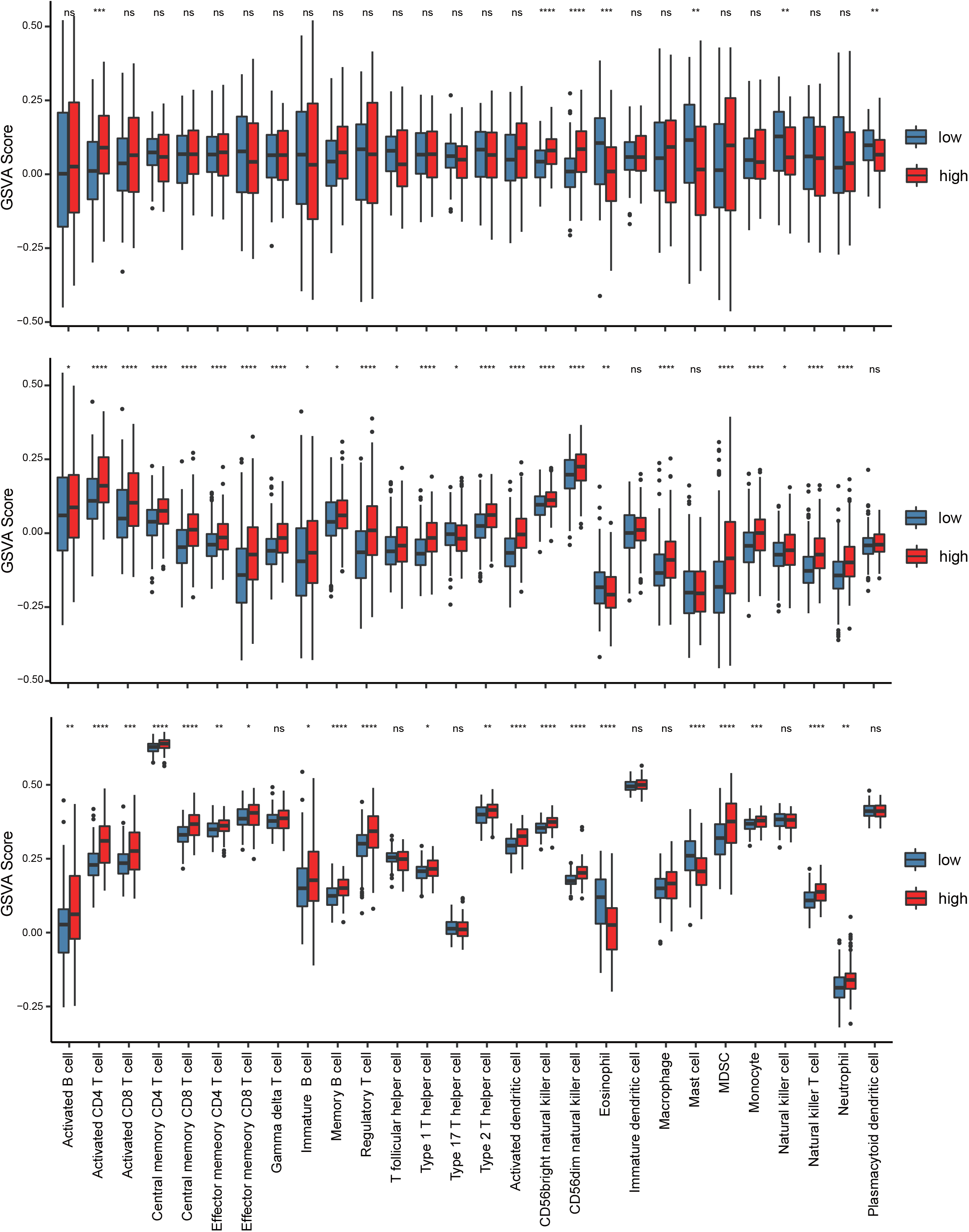
Stratified analysis of the Immune gene set in the external database. Comparative analysis of Immune gene set GSVA score according to the median R-index stratification in OncoSG, TCGA_LUAD, and GSE31210 database.

---

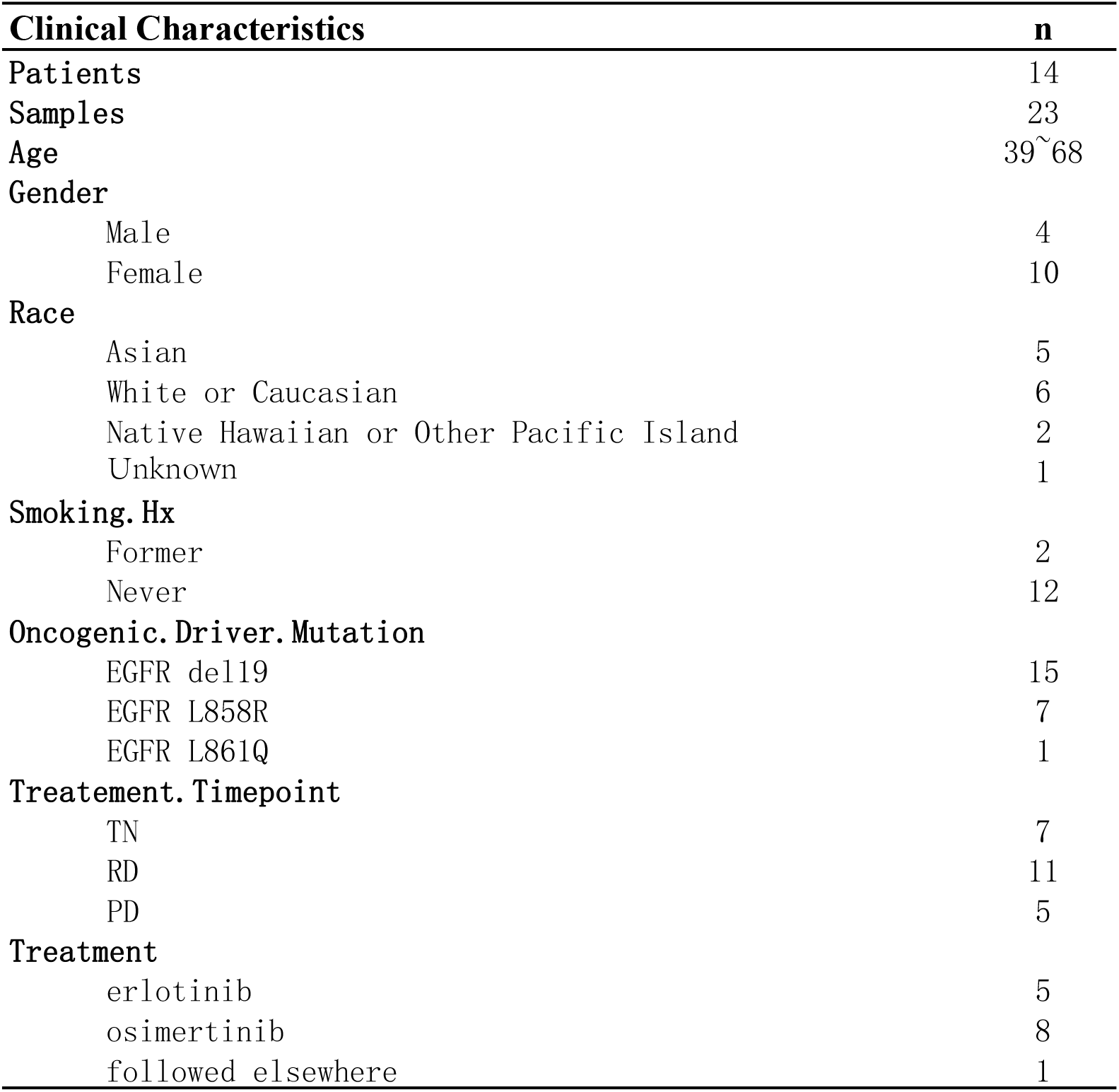

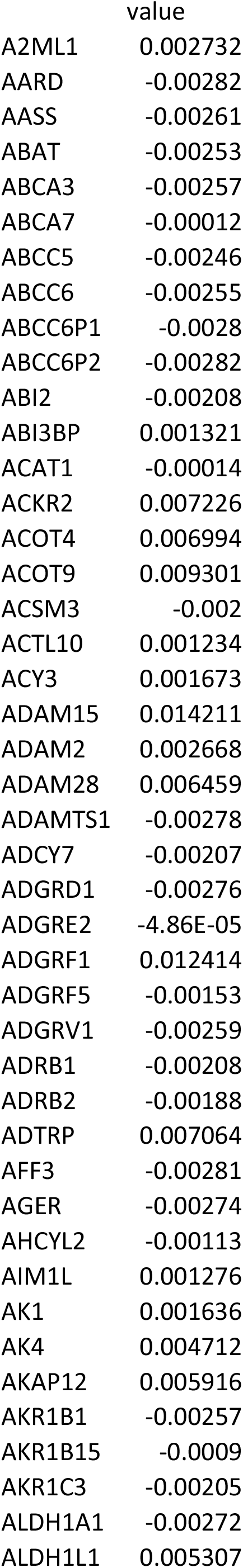
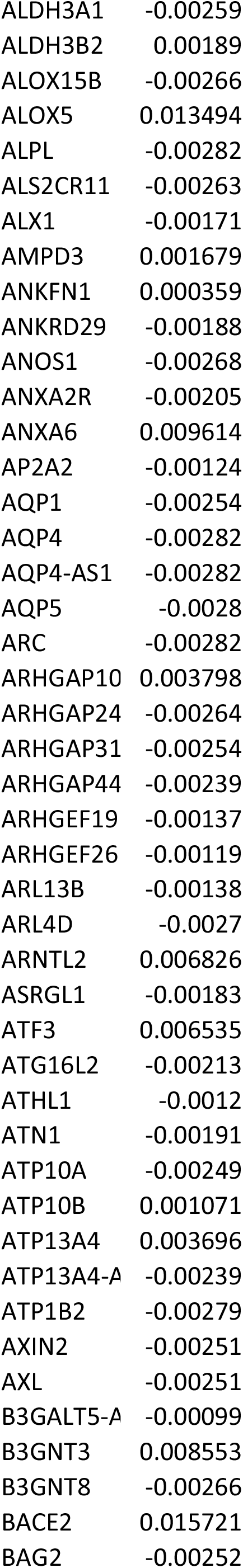
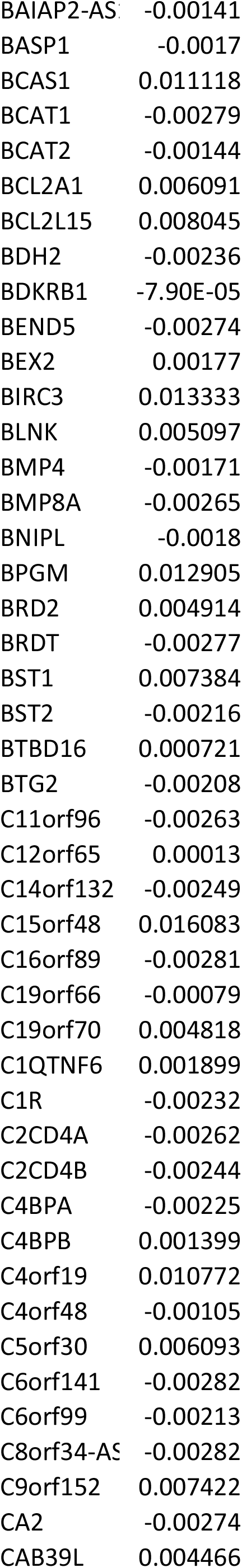
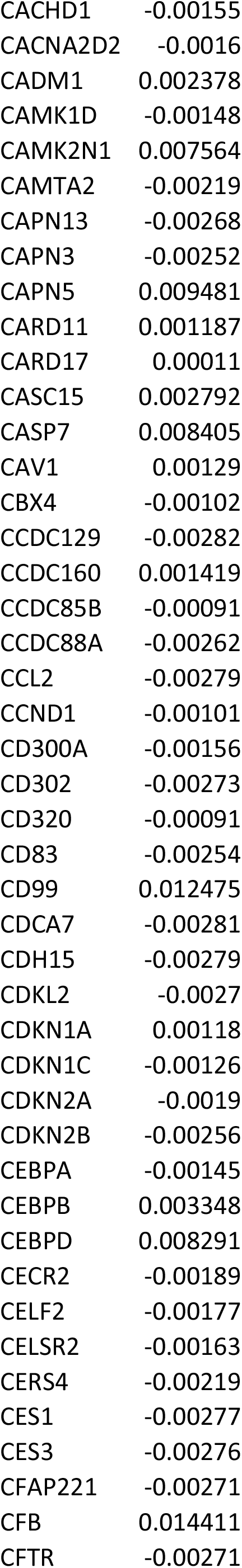
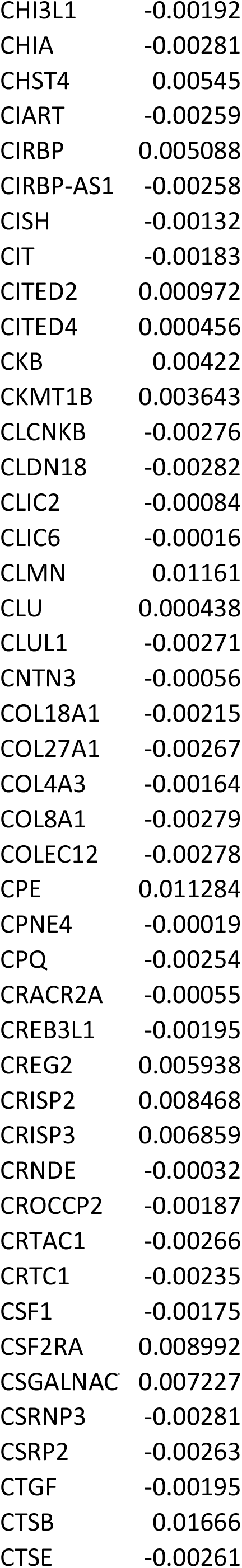
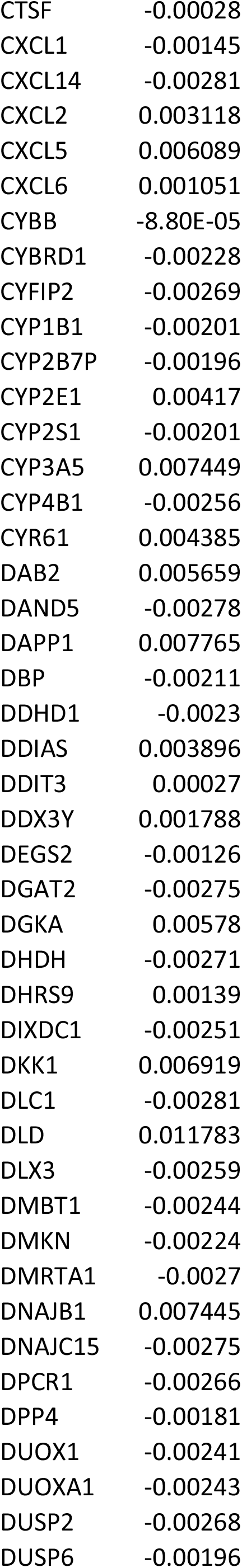
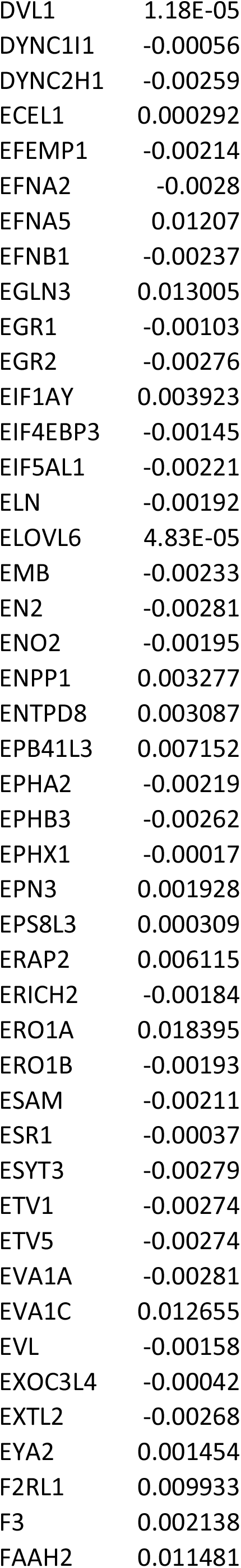
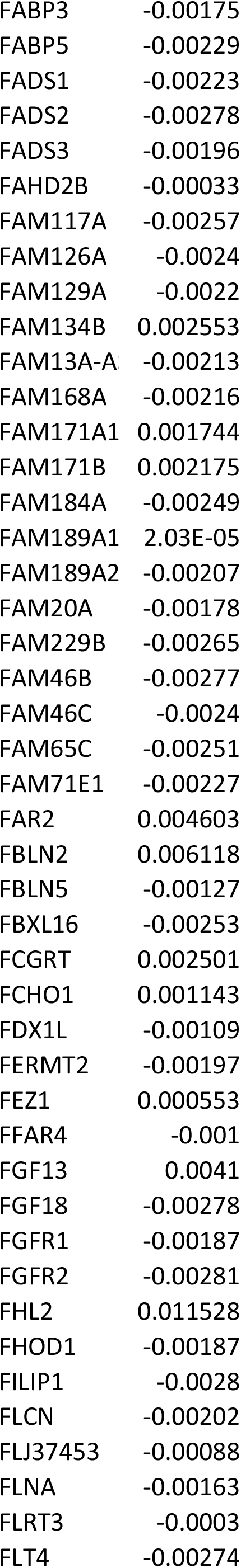
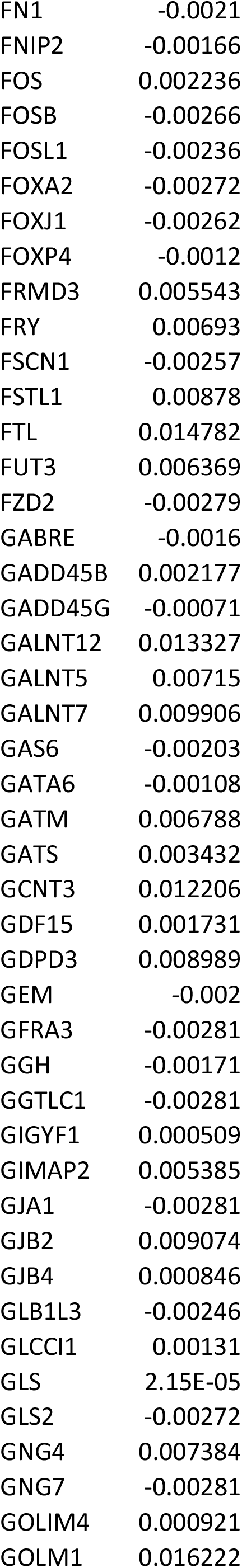
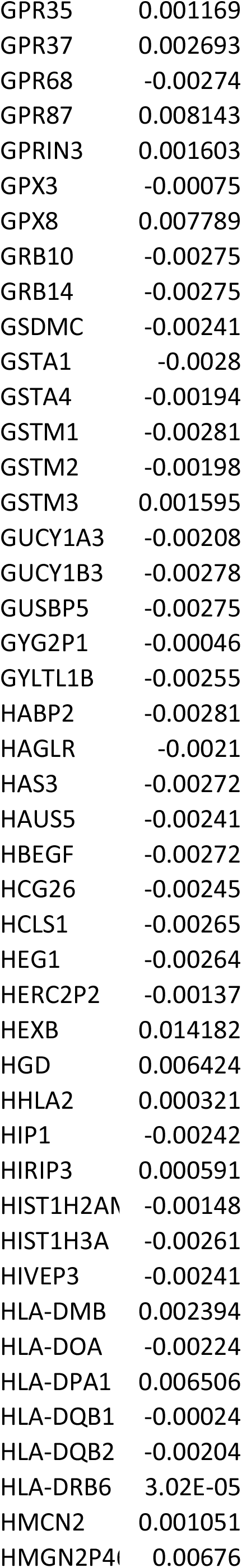
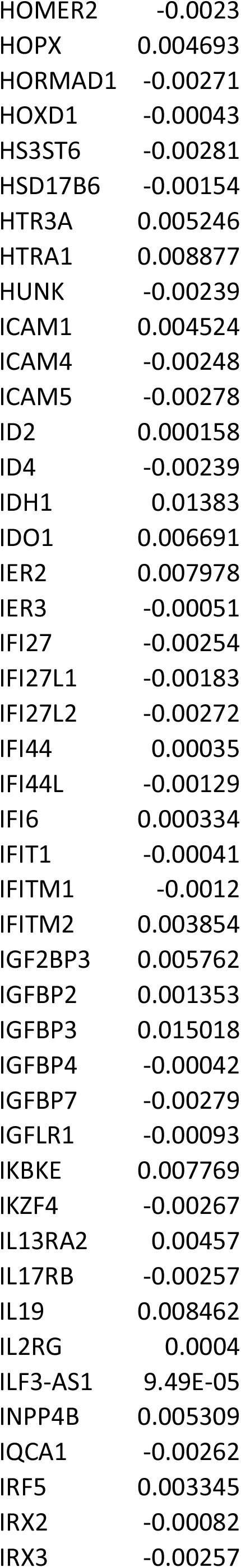
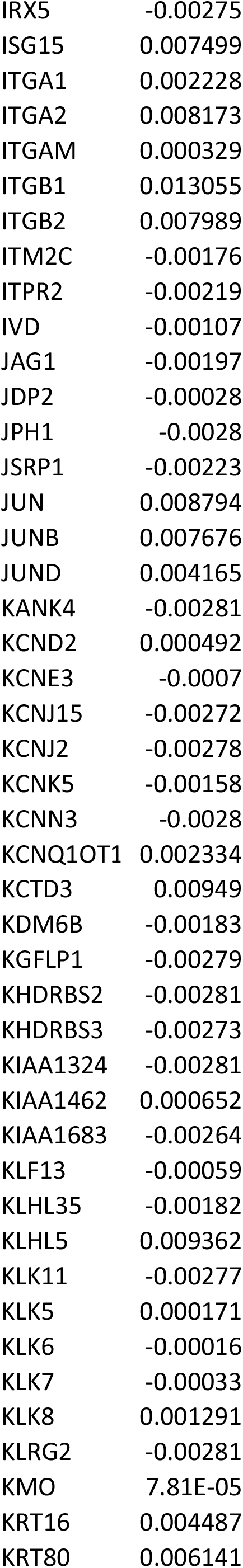
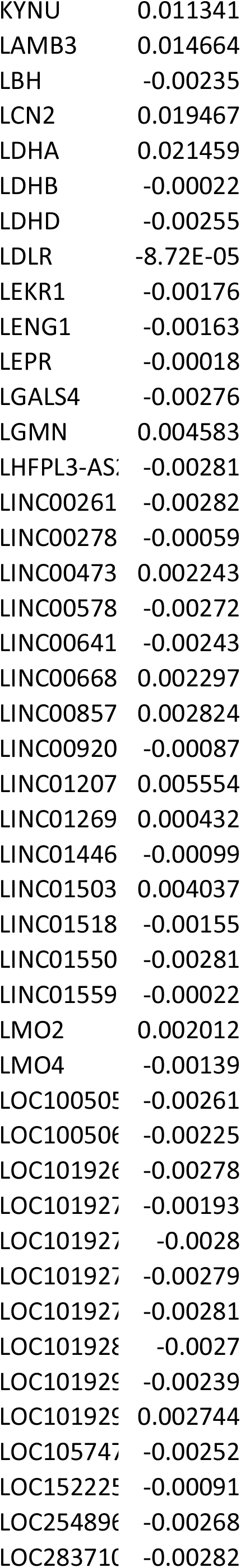
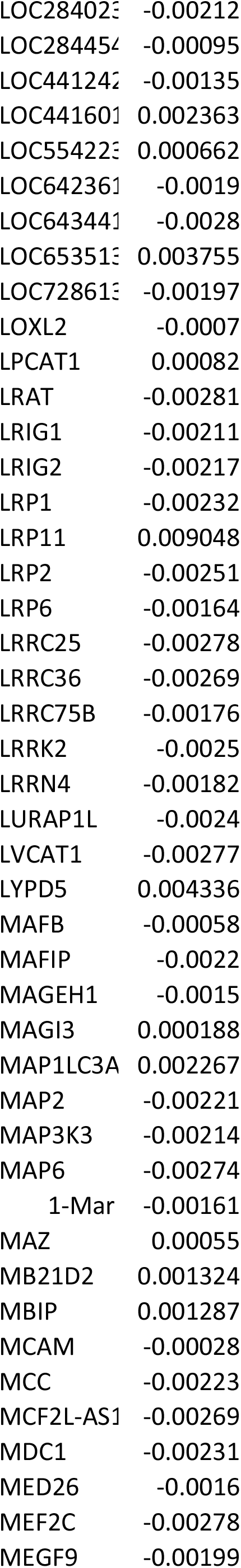
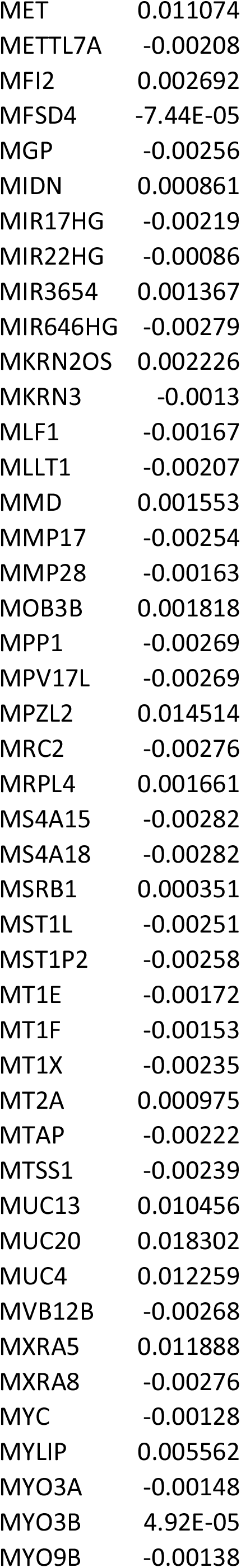
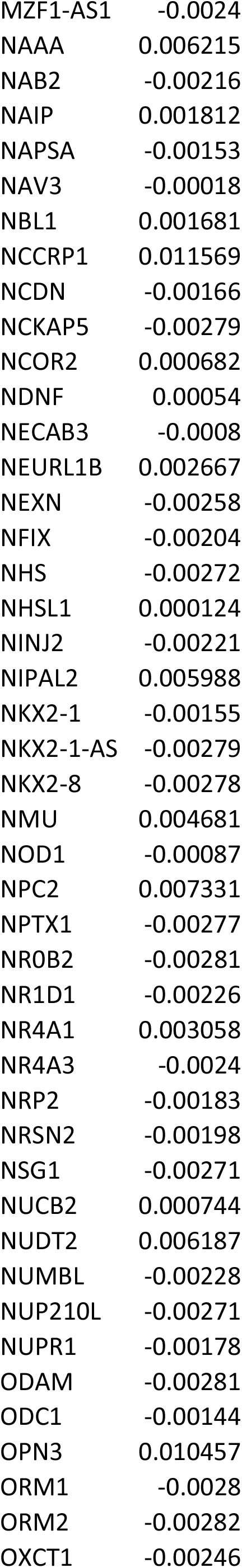
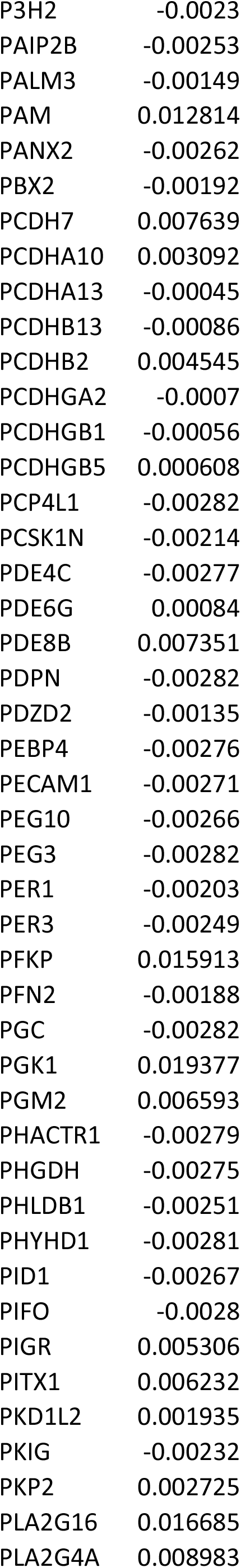
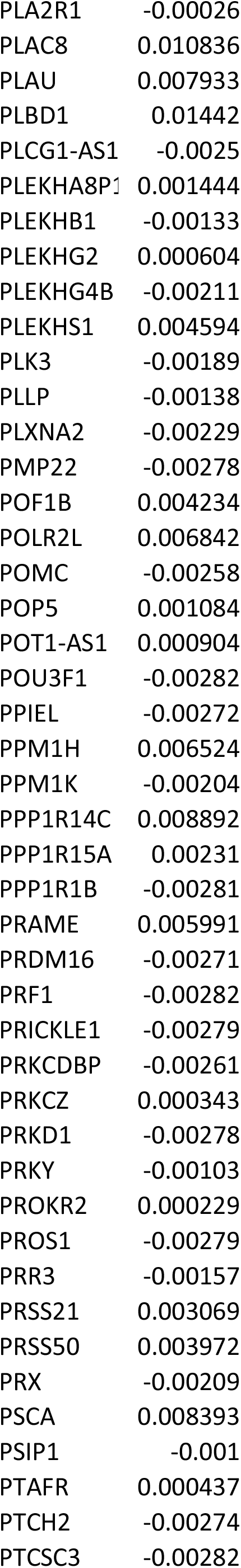
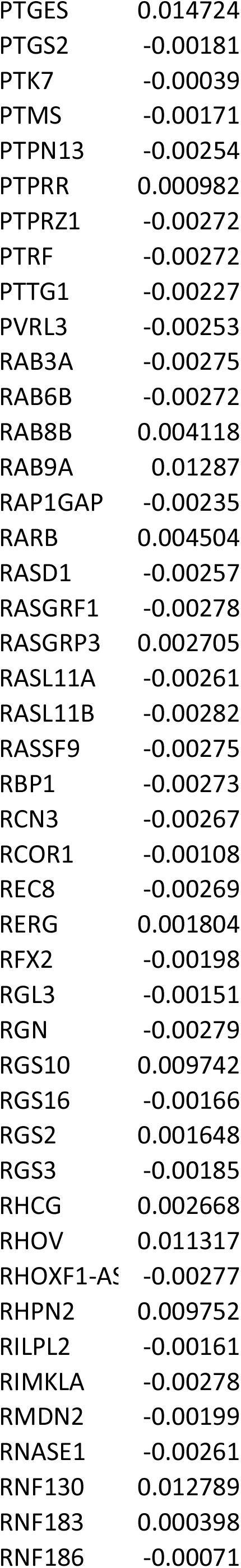
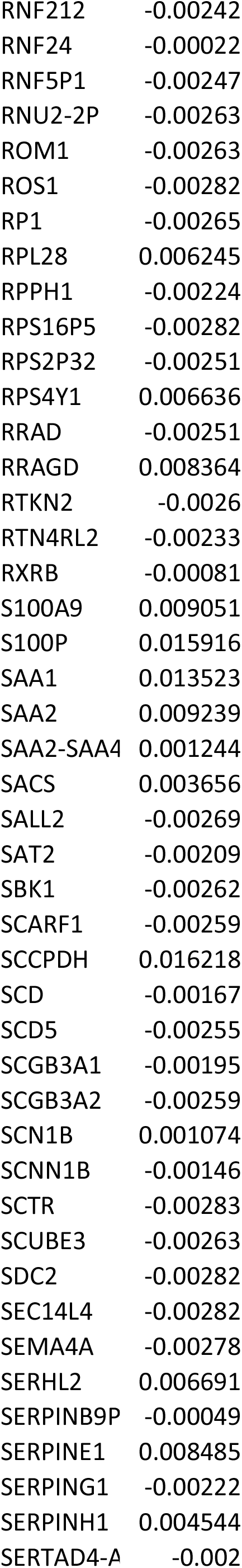
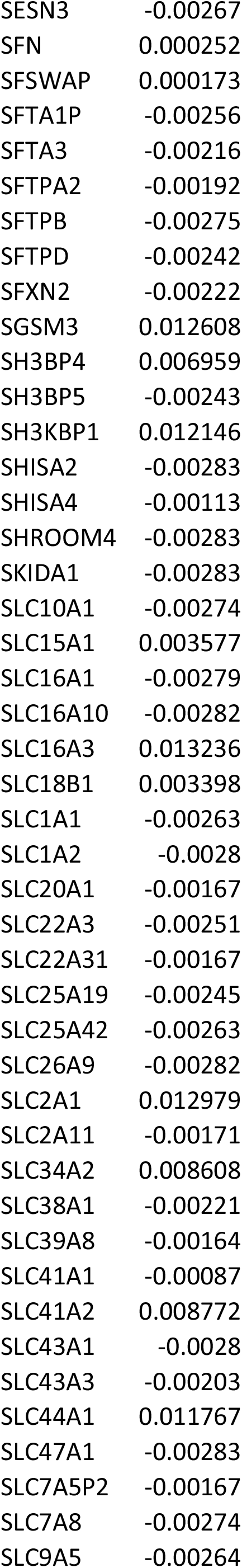
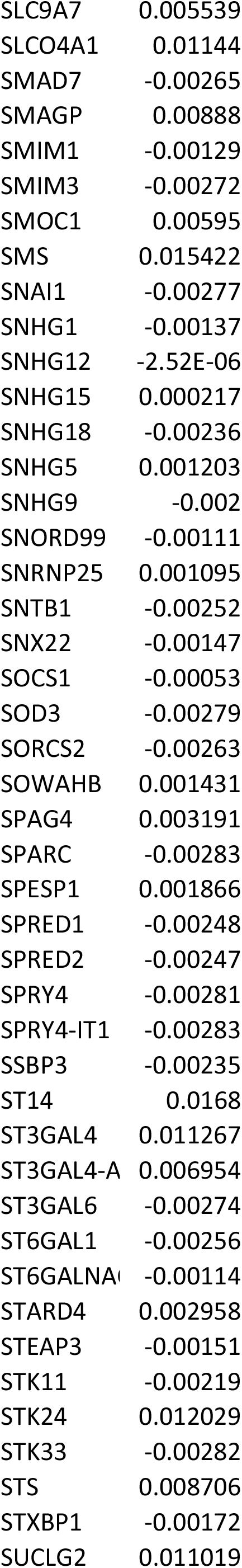
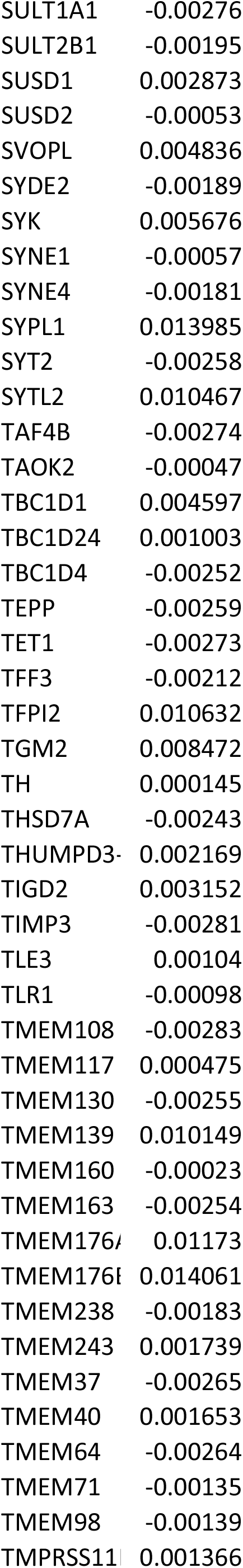
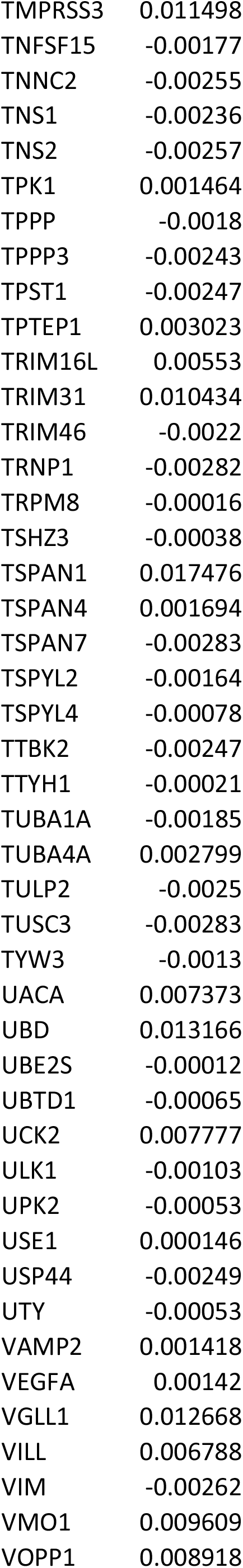
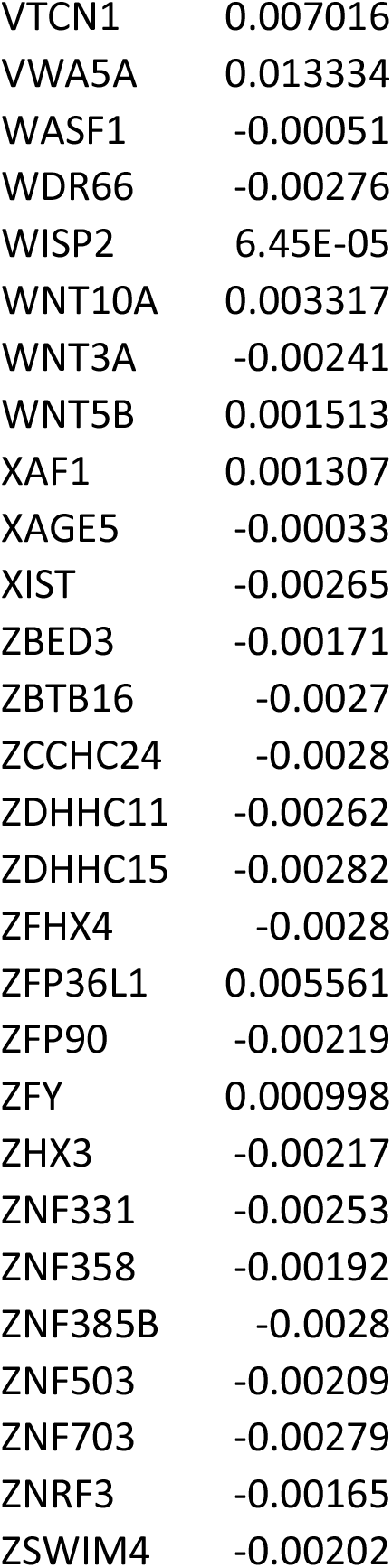

## REFERENCES

(2014) Comprehensive molecular profiling of lung adenocarcinoma. Nature 511: 543–50

Aissa AF, Islam A, Ariss MM, Go CC, Rader AE, Conrardy RD, Gajda AM, Rubio-Perez C, Valyi-Nagy K, Pasquinelli M, Feldman LE, Green SJ, Lopez-Bigas N, Frolov MV, Benevolenskaya EV (2021) Single-cell transcriptional changes associated with drug tolerance and response to combination therapies in cancer. Nature communications 12: 1628

Aktipis CA, Kwan VS, Johnson KA, Neuberg SL, Maley CC (2011) Overlooking evolution: a systematic analysis of cancer relapse and therapeutic resistance research. PloS one 6: e26100

Arcila ME, Oxnard GR, Nafa K, Riely GJ, Solomon SB, Zakowski MF, Kris MG, Pao W, Miller VA, Ladanyi M (2011) Rebiopsy of lung cancer patients with acquired resistance to EGFR inhibitors and enhanced detection of the T790M mutation using a locked nucleic acid-based assay. Clinical cancer research : an official journal of the American Association for Cancer Research 17: 1169–80

Bar J, Onn A (2012) Overcoming molecular mechanisms of resistance to first-generation epidermal growth factor receptor tyrosine kinase inhibitors. Clinical lung cancer 13: 267–79

Barretina J, Caponigro G, Stransky N, Venkatesan K, Margolin AA, Kim S, Wilson CJ, Lehár J, Kryukov GV, Sonkin D, Reddy A, Liu M, Murray L, Berger MF, Monahan JE, Morais P, Meltzer J, Korejwa A, Jané-Valbuena J, Mapa FA et al. (2012) The Cancer Cell Line Encyclopedia enables predictive modelling of anticancer drug sensitivity. Nature 483: 603–7

Barretina J, Caponigro G, Stransky N, Venkatesan K, Margolin AA, Kim S, Wilson CJ, Lehár J, Kryukov GV, Sonkin D, Reddy A, Liu M, Murray L, Berger MF, Monahan JE, Morais P, Meltzer J, Korejwa A, Jané-Valbuena J, Mapa FA et al. (2019) Addendum: The Cancer Cell Line Encyclopedia enables predictive modelling of anticancer drug sensitivity. Nature 565: E5–e6

Bronte V, Brandau S, Chen SH, Colombo MP, Frey AB, Greten TF, Mandruzzato S, Murray PJ, Ochoa A, Ostrand-Rosenberg S, Rodriguez PC, Sica A, Umansky V, Vonderheide RH, Gabrilovich DI (2016) Recommendations for myeloid-derived suppressor cell nomenclature and characterization standards. Nature communications 7: 12150

Butler A, Hoffman P, Smibert P, Papalexi E, Satija R (2018) Integrating single-cell transcriptomic data across different conditions, technologies, and species. Nature biotechnology 36: 411–420

Charoentong P, Finotello F, Angelova M, Mayer C, Efremova M, Rieder D, Hackl H, Trajanoski Z (2017) Pan-cancer Immunogenomic Analyses Reveal Genotype-Immunophenotype Relationships and Predictors of Response to Checkpoint Blockade. Cell reports 18: 248–262

Chen J, Yang H, Teo ASM, Amer LB, Sherbaf FG, Tan CQ, Alvarez JJS, Lu B, Lim JQ, Takano A, Nahar R, Lee YY, Phua CZJ, Chua KP, Suteja L, Chen PJ, Chang MM, Koh TPT, Ong BH, Anantham D et al. (2020) Genomic landscape of lung adenocarcinoma in East Asians. Nature genetics 52: 177–186

Chen S, Akdemir I, Fan J, Linden J, Zhang B, Cekic C (2020) The Expression of Adenosine A2B Receptor on Antigen-Presenting Cells Suppresses CD8(+) T-cell Responses and Promotes Tumor Growth. Cancer immunology research 8: 1064–1074

Chmielecki J, Foo J, Oxnard GR, Hutchinson K, Ohashi K, Somwar R, Wang L, Amato KR, Arcila M, Sos ML, Socci ND, Viale A, de Stanchina E, Ginsberg MS, Thomas RK, Kris MG, Inoue A, Ladanyi M, Miller VA, Michor F et al. (2011) Optimization of dosing for EGFR-mutant non-small cell lung cancer with evolutionary cancer modeling. Science translational medicine 3: 90ra59

Chow MT, Ozga AJ, Servis RL, Frederick DT, Lo JA, Fisher DE, Freeman GJ, Boland GM, Luster AD (2019) Intratumoral Activity of the CXCR3 Chemokine System Is Required for the Efficacy of Anti-PD-1 Therapy. Immunity 50: 1498–1512.e5

Domagala M, Laplagne C, Leveque E, Laurent C, Fournié JJ, Espinosa E, Poupot M (2021) Cancer Cells Resistance Shaping by Tumor Infiltrating Myeloid Cells. Cancers 13

Fukuoka M, Wu YL, Thongprasert S, Sunpaweravong P, Leong SS, Sriuranpong V, Chao TY, Nakagawa K, Chu DT, Saijo N, Duffield EL, Rukazenkov Y, Speake G, Jiang H, Armour AA, To KF, Yang JC, Mok TS (2011) Biomarker analyses and final overall survival results from a phase III, randomized, open-label, first-line study of gefitinib versus carboplatin/paclitaxel in clinically selected patients with advanced non-small-cell lung cancer in Asia (IPASS). Journal of clinical oncology : official journal of the American Society of Clinical Oncology 29: 2866–74

Gatenby RA, Brown JS (2020) Integrating evolutionary dynamics into cancer therapy. Nature reviews Clinical oncology 17: 675–686

Gatenby RA, Cunningham JJ, Brown JS (2014) Evolutionary triage governs fitness in driver and passenger mutations and suggests targeting never mutations. Nature communications 5: 5499

Hänzelmann S, Castelo R, Guinney J (2013) GSVA: gene set variation analysis for microarray and RNA-seq data. BMC bioinformatics 14: 7

He Y, Jia K, Dziadziuszko R, Zhao S, Zhang X, Deng J, Wang H, Hirsch FR, Zhou C (2019) Galectin-9 in non-small cell lung cancer. Lung cancer (Amsterdam, Netherlands) 136: 80–85

Hirschhaeuser F, Sattler UG, Mueller-Klieser W (2011) Lactate: a metabolic key player in cancer. Cancer research 71: 6921–5

Hoadley KA, Yau C, Hinoue T, Wolf DM, Lazar AJ, Drill E, Shen R, Taylor AM, Cherniack AD, Thorsson V, Akbani R, Bowlby R, Wong CK, Wiznerowicz M, Sanchez-Vega F, Robertson AG, Schneider BG, Lawrence MS, Noushmehr H, Malta TM et al. (2018) Cell-of-Origin Patterns Dominate the Molecular Classification of 10,000 Tumors from 33 Types of Cancer. Cell 173: 291–304.e6

Isomoto K, Haratani K, Hayashi H, Shimizu S, Tomida S, Niwa T, Yokoyama T, Fukuda Y, Chiba Y, Kato R, Tanizaki J, Tanaka K, Takeda M, Ogura T, Ishida T, Ito A, Nakagawa K (2020) Impact of EGFR-TKI Treatment on the Tumor Immune Microenvironment in EGFR Mutation-Positive Non-Small Cell Lung Cancer. Clinical cancer research : an official journal of the American Association for Cancer Research 26: 2037–2046

Jiang P, Gu S, Pan D, Fu J, Sahu A, Hu X, Li Z, Traugh N, Bu X, Li B, Liu J, Freeman GJ, Brown MA, Wucherpfennig KW, Liu XS (2018) Signatures of T cell dysfunction and exclusion predict cancer immunotherapy response. Nature medicine 24: 1550–1558

Johnson KE, Howard G, Mo W, Strasser MK, Lima E, Huang S, Brock A (2019) Cancer cell population growth kinetics at low densities deviate from the exponential growth model and suggest an Allee effect. PLoS biology 17: e3000399

Joshi K, de Massy MR, Ismail M, Reading JL, Uddin I, Woolston A, Hatipoglu E, Oakes T, Rosenthal R, Peacock T, Ronel T, Noursadeghi M, Turati V, Furness AJS, Georgiou A, Wong YNS, Ben Aissa A, Sunderland MW, Jamal-Hanjani M, Veeriah S et al. (2019) Spatial heterogeneity of the T cell receptor repertoire reflects the mutational landscape in lung cancer. Nature medicine 25: 1549–1559

Kim SM, Yun MR, Hong YK, Solca F, Kim JH, Kim HJ, Cho BC (2013) Glycolysis inhibition sensitizes non-small cell lung cancer with T790M mutation to irreversible EGFR inhibitors via translational suppression of Mcl-1 by AMPK activation. Molecular cancer therapeutics 12: 2145–56

Kita K, Fukuda K, Takahashi H, Tanimoto A, Nishiyama A, Arai S, Takeuchi S, Yamashita K, Ohtsubo K, Otani S, Yanagimura N, Suzuki C, Ikeda H, Tamura M, Matsumoto I, Yano S (2019) Patient-derived xenograft models of non-small cell lung cancer for evaluating targeted drug sensitivity and resistance. Cancer science 110: 3215–3224

Kitajima S, Asahina H, Chen T, Guo S, Quiceno LG, Cavanaugh JD, Merlino AA, Tange S, Terai H, Kim JW, Wang X, Zhou S, Xu M, Wang S, Zhu Z, Thai TC, Takahashi C, Wang Y, Neve R, Stinson S et al. (2018) Overcoming Resistance to Dual Innate Immune and MEK Inhibition Downstream of KRAS. Cancer cell 34: 439–452.e6

Kobayashi S, Boggon TJ, Dayaram T, Jänne PA, Kocher O, Meyerson M, Johnson BE, Eck MJ, Tenen DG, Halmos B (2005) EGFR mutation and resistance of non-small-cell lung cancer to gefitinib. The New England journal of medicine 352: 786–92

Korotkevich G, Sukhov V, Budin N, Shpak B, Artyomov MN, Sergushichev A (2021) Fast gene set enrichment analysis. bioRxiv: 060012

Lee SM, Park CM, Lee KH, Bahn YE, Kim JI, Goo JM (2014) C-arm cone-beam CT-guided percutaneous transthoracic needle biopsy of lung nodules: clinical experience in 1108 patients. Radiology 271: 291–300

Li R, Salehi-Rad R, Crosson W, Momcilovic M, Lim RJ, Ong SL, Huang ZL, Zhang T, Abascal J, Dumitras C, Jing Z, Park SJ, Krysan K, Shackelford DB, Tran LM, Liu B, Dubinett SM (2021) Inhibition of Granulocytic Myeloid-Derived Suppressor Cells Overcomes Resistance to Immune Checkpoint Inhibition in LKB1-deficient Non-Small Cell Lung Cancer. Cancer research

Liberzon A, Birger C, Thorvaldsdóttir H, Ghandi M, Mesirov JP, Tamayo P (2015) The Molecular Signatures Database (MSigDB) hallmark gene set collection. Cell systems 1: 417–425

Lito P, Rosen N, Solit DB (2013) Tumor adaptation and resistance to RAF inhibitors. Nature medicine 19: 1401–9

Liu H, Kuang X, Zhang Y, Ye Y, Li J, Liang L, Xie Z, Weng L, Guo J, Li H, Ma F, Chen X, Zhao S, Su J, Yang N, Fang F, Xie Y, Tao J, Zhang J, Chen M et al. (2020) ADORA1 Inhibition Promotes Tumor Immune Evasion by Regulating the ATF3-PD-L1 Axis. Cancer cell 37: 324–339.e8

Lynch TJ, Bell DW, Sordella R, Gurubhagavatula S, Okimoto RA, Brannigan BW, Harris PL, Haserlat SM, Supko JG, Haluska FG, Louis DN, Christiani DC, Settleman J, Haber DA (2004) Activating mutations in the epidermal growth factor receptor underlying responsiveness of non-small-cell lung cancer to gefitinib. The New England journal of medicine 350: 2129–39

Malta TM, Sokolov A, Gentles AJ, Burzykowski T, Poisson L, Weinstein JN, Kamińska B, Huelsken J, Omberg L, Gevaert O, Colaprico A, Czerwińska P, Mazurek S, Mishra L, Heyn H, Krasnitz A, Godwin AK, Lazar AJ, Stuart JM, Hoadley KA et al. (2018) Machine Learning Identifies Stemness Features Associated with Oncogenic Dedifferentiation. Cell 173: 338–354.e15

Maynard A, McCoach CE, Rotow JK, Harris L, Haderk F, Kerr DL, Yu EA, Schenk EL, Tan W, Zee A, Tan M, Gui P, Lea T, Wu W, Urisman A, Jones K, Sit R, Kolli PK, Seeley E, Gesthalter Y et al. (2020) Therapy-Induced Evolution of Human Lung Cancer Revealed by Single-Cell RNA Sequencing. Cell 182: 1232–1251.e22

Moesta AK, Li XY, Smyth MJ (2020) Targeting CD39 in cancer. Nature reviews Immunology 20: 739–755

Molina JR, Yang P, Cassivi SD, Schild SE, Adjei AA (2008) Non-small cell lung cancer: epidemiology, risk factors, treatment, and survivorship. Mayo Clinic proceedings 83: 584–94

Oh IJ, Ban HJ, Kim KS, Kim YC (2012) Retreatment of gefitinib in patients with non-small-cell lung cancer who previously controlled to gefitinib: a single-arm, open-label, phase II study. Lung cancer (Amsterdam, Netherlands) 77: 121–7

Ohashi K, Maruvka YE, Michor F, Pao W (2013) Epidermal growth factor receptor tyrosine kinase inhibitor-resistant disease. Journal of clinical oncology : official journal of the American Society of Clinical Oncology 31: 1070–80

Okayama H, Kohno T, Ishii Y, Shimada Y, Shiraishi K, Iwakawa R, Furuta K, Tsuta K, Shibata T, Yamamoto S, Watanabe S, Sakamoto H, Kumamoto K, Takenoshita S, Gotoh N, Mizuno H, Sarai A, Kawano S, Yamaguchi R, Miyano S et al. (2012) Identification of genes upregulated in ALK-positive and EGFR/KRAS/ALK-negative lung adenocarcinomas. Cancer research 72: 100–11

Paez JG, Jänne PA, Lee JC, Tracy S, Greulich H, Gabriel S, Herman P, Kaye FJ, Lindeman N, Boggon TJ, Naoki K, Sasaki H, Fujii Y, Eck MJ, Sellers WR, Johnson BE, Meyerson M (2004) EGFR mutations in lung cancer: correlation with clinical response to gefitinib therapy. Science (New York, N) 304: 1497–500

Peng S, Wang R, Zhang X, Ma Y, Zhong L, Li K, Nishiyama A, Arai S, Yano S, Wang W (2019) EGFR-TKI resistance promotes immune escape in lung cancer via increased PD-L1 expression. Molecular cancer 18: 165

Planchard D, Popat S, Kerr K, Novello S, Smit EF, Faivre-Finn C, Mok TS, Reck M, Van Schil PE, Hellmann MD, Peters S (2018) Metastatic non-small cell lung cancer: ESMO Clinical Practice Guidelines for diagnosis, treatment and follow-up. Annals of oncology : official journal of the European Society for Medical Oncology 29: iv192–iv237

Qiu X, Mao Q, Tang Y, Wang L, Chawla R, Pliner HA, Trapnell C (2017) Reversed graph embedding resolves complex single-cell trajectories. Nature methods 14: 979–982

Racker E, Resnick RJ, Feldman R (1985) Glycolysis and methylaminoisobutyrate uptake in rat-1 cells transfected with ras or myc oncogenes. Proceedings of the National Academy of Sciences of the United States of America 82: 3535–8

Rosell R, Moran T, Queralt C, Porta R, Cardenal F, Camps C, Majem M, Lopez-Vivanco G, Isla D, Provencio M, Insa A, Massuti B, Gonzalez-Larriba JL, Paz-Ares L, Bover I, Garcia-Campelo R, Moreno MA, Catot S, Rolfo C, Reguart N et al. (2009) Screening for epidermal growth factor receptor mutations in lung cancer. The New England journal of medicine 361: 958–67

Scapini P, Cassatella MA (2014) Social networking of human neutrophils within the immune system. Blood 124: 710–9

Sequist LV, Waltman BA, Dias-Santagata D, Digumarthy S, Turke AB, Fidias P, Bergethon K, Shaw AT, Gettinger S, Cosper AK, Akhavanfard S, Heist RS, Temel J, Christensen JG, Wain JC, Lynch TJ, Vernovsky K, Mark EJ, Lanuti M, Iafrate AJ et al. (2011) Genotypic and histological evolution of lung cancers acquiring resistance to EGFR inhibitors. Science translational medicine 3: 75ra26

Sharma SV, Lee DY, Li B, Quinlan MP, Takahashi F, Maheswaran S, McDermott U, Azizian N, Zou L, Fischbach MA, Wong KK, Brandstetter K, Wittner B, Ramaswamy S, Classon M, Settleman J (2010) A chromatin-mediated reversible drug-tolerant state in cancer cell subpopulations. Cell 141: 69–80

Shaul ME, Fridlender ZG (2019) Tumour-associated neutrophils in patients with cancer. Nature reviews Clinical oncology 16: 601–620

Shen F, Chu S, Bence AK, Bailey B, Xue X, Erickson PA, Montrose MH, Beck WT, Erickson LC (2008) Quantitation of doxorubicin uptake, efflux, and modulation of multidrug resistance (MDR) in MDR human cancer cells. The Journal of pharmacology and experimental therapeutics 324: 95–102

Siegel RL, Miller KD, Fuchs HE, Jemal A (2021) Cancer Statistics, 2021. CA: a cancer journal for clinicians 71: 7–33

Sokolov A, Paull EO, Stuart JM (2016) ONE-CLASS DETECTION OF CELL STATES IN TUMOR SUBTYPES. Pacific Symposium on Biocomputing Pacific Symposium on Biocomputing 21: 405–16

Song T, Yu W, Wu SX (2014) Subsequent treatment choices for patients with acquired resistance to EGFR-TKIs in non-small cell lung cancer: restore after a drug holiday or switch to another EGFR-TKI? Asian Pacific journal of cancer prevention : APJCP 15: 205–13

Sun Y, Wu L, Zhong Y, Zhou K, Hou Y, Wang Z, Zhang Z, Xie J, Wang C, Chen D, Huang Y, Wei X, Shi Y, Zhao Z, Li Y, Guo Z, Yu Q, Xu L, Volpe G, Qiu S et al. (2021) Single-cell landscape of the ecosystem in early-relapse hepatocellular carcinoma. Cell 184: 404–421.e16

Suzuki S, Okada M, Takeda H, Kuramoto K, Sanomachi T, Togashi K, Seino S, Yamamoto M, Yoshioka T, Kitanaka C (2018) Involvement of GLUT1-mediated glucose transport and metabolism in gefitinib resistance of non-small-cell lung cancer cells. Oncotarget 9: 32667–32679

Tamada M, Nagano O, Tateyama S, Ohmura M, Yae T, Ishimoto T, Sugihara E, Onishi N, Yamamoto T, Yanagawa H, Suematsu M, Saya H (2012) Modulation of glucose metabolism by CD44 contributes to antioxidant status and drug resistance in cancer cells. Cancer research 72: 1438–48

Tomiyama A, Serizawa S, Tachibana K, Sakurada K, Samejima H, Kuchino Y, Kitanaka C (2006) Critical role for mitochondrial oxidative phosphorylation in the activation of tumor suppressors Bax and Bak. Journal of the National Cancer Institute 98: 1462–73

Vander Heiden MG, Cantley LC, Thompson CB (2009) Understanding the Warburg effect: the metabolic requirements of cell proliferation. Science (New York, NY) 324: 1029–33

Walther V, Hiley CT, Shibata D, Swanton C, Turner PE, Maley CC (2015) Can oncology recapitulate paleontology? Lessons from species extinctions. Nature reviews Clinical oncology 12: 273–85

Warburg O (1956) On the origin of cancer cells. Science (New York, NY) 123: 309–14

Watanabe S, Tanaka J, Ota T, Kondo R, Tanaka H, Kagamu H, Ichikawa K, Koshio J, Baba J, Miyabayashi T, Narita I, Yoshizawa H (2011) Clinical responses to EGFR-tyrosine kinase inhibitor retreatment in non-small cell lung cancer patients who benefited from prior effective gefitinib therapy: a retrospective analysis. BMC cancer 11: 1

Weber R, Fleming V, Hu X, Nagibin V, Groth C, Altevogt P, Utikal J, Umansky V (2018) Myeloid-Derived Suppressor Cells Hinder the Anti-Cancer Activity of Immune Checkpoint Inhibitors. Frontiers in immunology 9: 1310

Wu SG, Shih JY (2018) Management of acquired resistance to EGFR TKI-targeted therapy in advanced non-small cell lung cancer. Molecular cancer 17: 38

Xie H, Hanai J, Ren JG, Kats L, Burgess K, Bhargava P, Signoretti S, Billiard J, Duffy KJ, Grant A, Wang X, Lorkiewicz PK, Schatzman S, Bousamra M, 2nd, Lane AN, Higashi RM, Fan TW, Pandolfi PP, Sukhatme VP, Seth P (2014) Targeting lactate dehydrogenase--a inhibits tumorigenesis and tumor progression in mouse models of lung cancer and impacts tumor-initiating cells. Cell metabolism 19: 795–809

Yamaguchi O, Kaira K, Mouri A, Shiono A, Hashimoto K, Miura Y, Nishihara F, Murayama Y, Kobayashi K, Kagamu H (2019) Re-challenge of afatinib after 1st generation EGFR-TKI failure in patients with previously treated non-small cell lung cancer harboring EGFR mutation. Cancer chemotherapy and pharmacology 83: 817–825

Yang W, Soares J, Greninger P, Edelman EJ, Lightfoot H, Forbes S, Bindal N, Beare D, Smith JA, Thompson IR, Ramaswamy S, Futreal PA, Haber DA, Stratton MR, Benes C, McDermott U, Garnett MJ (2013) Genomics of Drug Sensitivity in Cancer (GDSC): a resource for therapeutic biomarker discovery in cancer cells. Nucleic acids research 41: D955–61

Yu HA, Arcila ME, Rekhtman N, Sima CS, Zakowski MF, Pao W, Kris MG, Miller VA, Ladanyi M, Riely GJ (2013) Analysis of tumor specimens at the time of acquired resistance to EGFR-TKI therapy in 155 patients with EGFR-mutant lung cancers. Clinical cancer research : an official journal of the American Association for Cancer Research 19: 2240–7

Yuan S, Norgard RJ, Stanger BZ (2019) Cellular Plasticity in Cancer. Cancer discovery 9: 837–851

Zhang J, Cunningham JJ, Brown JS, Gatenby RA (2017) Integrating evolutionary dynamics into treatment of metastatic castrate-resistant prostate cancer. Nature communications 8: 1816

Zhang Z, Lee JC, Lin L, Olivas V, Au V, LaFramboise T, Abdel-Rahman M, Wang X, Levine AD, Rho JK, Choi YJ, Choi CM, Kim SW, Jang SJ, Park YS, Kim WS, Lee DH, Lee JS, Miller VA, Arcila M et al. (2012) Activation of the AXL kinase causes resistance to EGFR-targeted therapy in lung cancer. Nature genetics 44: 852–60

